# DNA Origami Tension Sensors (DOTS) to study T cell receptor mechanics at membrane junctions

**DOI:** 10.1101/2023.07.09.548279

**Authors:** Yuesong Hu, Yuxin Duan, Arventh Velusamy, Steven Narum, Jhordan Rogers, Khalid Salaita

**Affiliations:** Department of Chemistry, Emory University, Atlanta, GA, United States; Wallace H. Coulter Department of Biomedical Engineering, Georgia Institute of Technology and Emory University, Atlanta, GA, United States

## Abstract

The T cell receptor (TCR) is thought to be a mechanosensor, meaning that it transmits mechanical force to its antigen and leverages the force to amplify the specificity and magnitude of TCR signaling. The past decade has witnessed the development of molecular probes which have revealed many aspects of receptor mechanotransduction. However, most force probes are immobilized on hard substrates, thus failing to reveal mechanics in the physiological context of cell membranes. In this report, we developed DNA origami tension sensors (DOTS) which bear force sensors on a DNA origami breadboard and allow mapping of TCR mechanotransduction at dynamic intermembrane junctions. We demonstrate that TCR-antigen bonds experience 5-10 pN forces, and the mechanical events are dependent on cell state, antigen mobility, antigen potency, antigen height and F-actin activity. We tethered DOTS onto a microparticle to mechanically screen antigen in high throughput using flow cytometry. Finally, DOTS were anchored onto live B cell membranes thus producing the first quantification of TCR mechanics at authentic immune cell-cell junctions.

## Introduction

To protect against cancer and viral infections, the T cell receptor (TCR) actively scans the surface of target cells seeking to recognize abnormal protein fragments presented by the major histocompatibility complex, pMHC. Remarkably, the TCR exhibits an exceptional level of specificity and sensitivity towards antigens. Even the presence of just one or two abnormal pMHC molecules among approximately 100,000 normal pMHCs on a single cell, is sufficient to trigger T cell activation.^1^ The molecular mechanisms responsible for such a robust T cell response are still not fully understood. However, an emerging hypothesis suggests that TCR-pMHC bonds within the dynamic environment of cell-cell junctions experience mechanical forces and these forces can cause conformation changes in the TCR-pMHC complexes, thereby prolonging the lifetime and exposing kinase docking sites to facilitate the subsequent phosphorylation cascade necessary for TCR signaling.^2, 3, 4^Indeed, studies using single molecule force techniques have demonstrated that ∼10 piconewton (pN) forces is capable of triggering calcium flux, hallmark of T-cell activation.^5, 6^However, these forces were externally applied to the pMHC-TCR complexes leaving the question of whether the magnitude and duration of such forces are representative of the native immune junction. This important inquiry highlights a prevailing challenge in the field, namely the need to develop a tool to investigate the TCR mechanotransduction at intermembrane junctions.

In the past decade, our lab and colleagues developed a series of molecular tension sensors (MTS) to detect the forces transmitted to individual receptor-ligand bonds. Briefly, an elastic molecule (DNA hairpin, peptide, or polymer) is modified with a FRET pair and anchored to a surface at one terminus and presenting a ligand to bind a receptor of interest at its other terminus.^7, 8, 9^Cellular forces transmitted to the probe extend it and separate the FRET pair, leading to a fluorescence enhancement. With this sensor, we revealed that TCRs transmit 10-20 pN forces to antigens and these forces contribute to antigen discrimination, TCR proximal signaling, T cell activation, and cytotoxic degranulation.^10, 11, 12^However, in these studies, sensors were immobilized on a hard glass slide, restricting lateral motion, unlike the case in cell membranes. This type of immobilization inhibits TCR clustering and centralization, potentially resulting in an overestimation of the force magnitude. Several studies, including our own, used glass supported lipid bilayers (SLBs) to mimic the plasma membrane and inserted MTS to SLB to measure force at laterally fluid TCR-antigen bonds.^13, 14^However, at TCR clusters, MTS are likely placed within sufficient proximity that causes crosstalk between FRET pairs at adjacent sensors. This would lead to suppressed fluorescence emission, thereby compromising the reliability of the results. This inconsistency in fluorescence levels may provide an explanation for the discrepancies observed in the magnitudes of TCR forces previously reported by different research groups utilizing MTS-based approaches. Additionally, it is important to note that the SLBs used in these studies were created on planar glass substrates. Planar SLBs lack deformability and exhibit a flattened topology, thus resulting in a flat and contiguous T cell/SLB contact zone. In contrast, physiological T cell-target cell contacts are highly dynamic, characterized by sporadic and curved interactions, often involving microvilli or invadosome-like protrusions.^15, 16^Moreover, glass substrates are chemically and physically different from that of an antigen presenting cell. For example, the stiffness of glass is 10^9^-fold greater than that of the plasma membrane and prior studies have showed that T cell mechanotransduction is influenced by the stiffness of the surrounding matrix.^17, 18^

Herein we developed membrane-tethered DNA Origami Tension Sensors (DOTS) to address these limitations. DNA origami, due to its high programmability and functionality, has emerged as a versatile approach to fabricate nanodevices for spatial patterning, sensing and molecular manipulation and provides a powerful platform for force sensor design.^19, 20, 21^Specifically, we made a rectangular nanosheet origami carrying a DNA hairpin that is responsive to TCR forces. The dimensions of the origami set the minimum distance between adjacent hairpins at 40 nm thus fully suppressing the crosstalk between FRET pairs. With DOTS, we revealed that TCR-antigen bonds experience force exceeding 8.4 pN at fluid interfaces and these forces was generated by F-actin dependent cytoskeleton contraction and repulsion from large proteins at the immune synapse. This conclusion was validated using both ratiometric fluorescence intensity measurements as well fluorescence lifetime-based imaging. To best mimic the geometry of antigen presenting cells and reconstitute the three-dimensionality of immune synapse, DOTS were tethered to SLB functionalized microparticles which allowed for measuring TCR force in suspension and in a high-throughput manner using flow cytometry, thus offering a potential tool for mechanically based antigen screening. Finally, we anchored DOTS to live B-cell membranes and thus creating the first molecular device that allows for measuring force transmission at authentic immune cell-cell junctions. Taken together, DOTS represent a powerful tool to study immunoreceptor forces and the technique is expected to promote advances in the emerging field of immunomechanobiology.

## Results

### Design and characterization of DOTS

We designed a single layer DNA origami nanostructure with dimension of 40 x 80 nm, which was folded by hybridizing 84 single stranded DNA oligos (staples) to p7560 phage DNA scaffold (**Fig. 1a** and **Supplementary Fig. 3**). A DNA hairpin sequence was incorporated into one staple and extended out of the origami to detect TCR forces. The integrity and structure of assembled DOTS was confirmed by gel electrophoresis and atomic force microscopy (AFM) (**Fig. 1b**). No degradation of origami was observed even after 1 hour incubation in cell imaging media at room temperature (RT) (**Supplementary Fig. 4**). The origami was anchored to the SLB surface by hybridizing eight single-stranded DNA overhangs located at the bottom of the origami to preinserted complementary cholesterol DNA strands on the SLB surface. Throughout this work, unless otherwise specified, we utilized a concentration of 5 nM origami to prepare the surfaces, resulting in a density of 370±13 molecules/*μ*m^2^ (**Supplementary Fig. 5**). Fluorescence Recover After Photobleaching (FRAP) measurements showed that DOTS were laterally mobile on DOPC SLB and exhibited a physiological diffusion coefficient of ∼0.04 *μ*m^2^/s. Note the fluidity of DOTS is similar to that of murine antigens on the plasma membrane. This is in contrast to previously reported fluid MTS which exhibited a supraphysiological fluidity that was 10-20 times higher (**Fig. 1c**).^13, 22^

**Figure 1.**
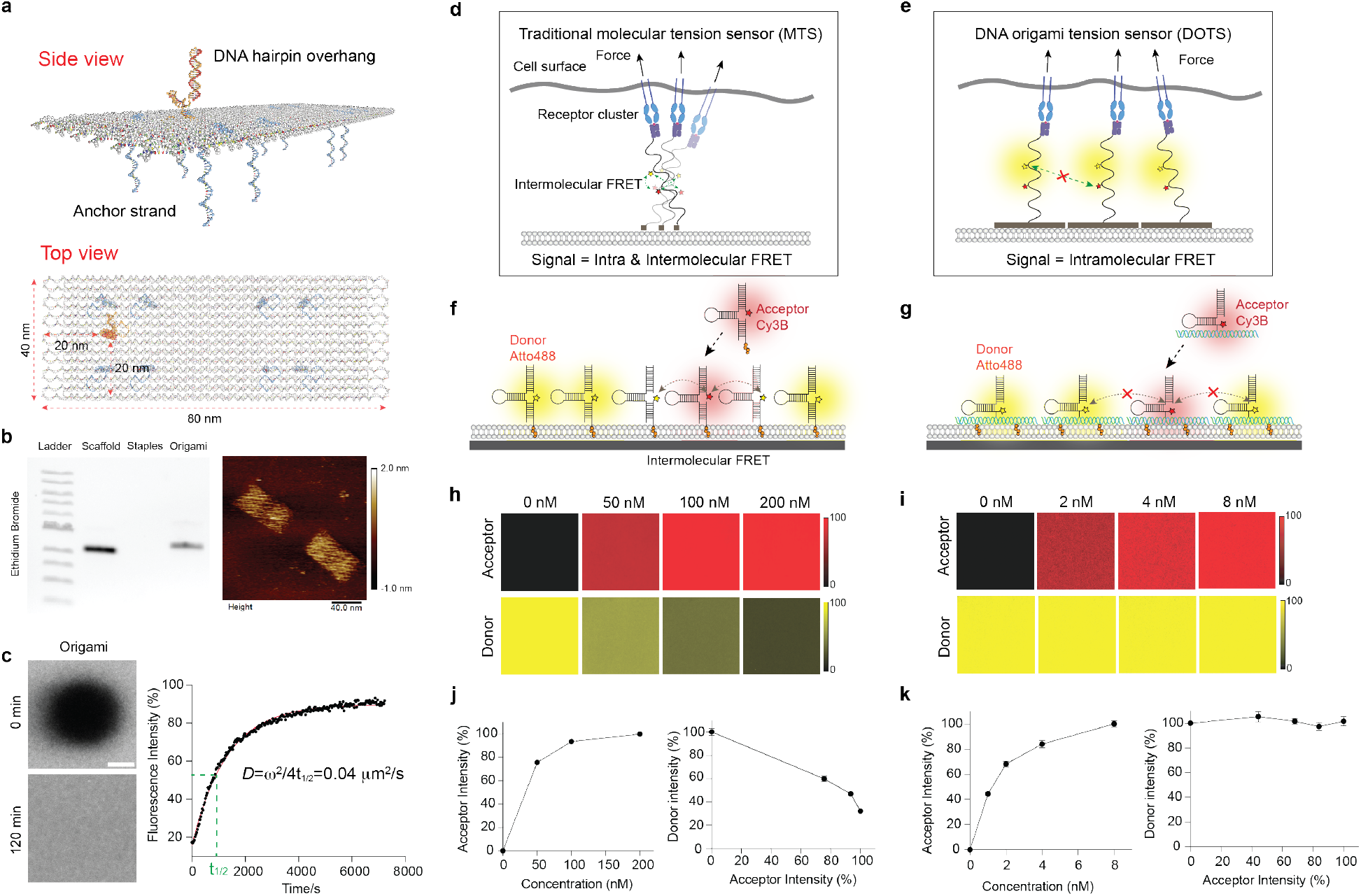
Characterization of DOTS. **a** Schematic of DOTS comprised of a rectangular nanosheet that presents a single hairpin on one face and eight anchor strands on the other face for anchoring DOTS to membrane. **b** Left: agarose gel electrophoresis showing the bands of DNA origami scaffold, staples, and annealed DOTS. Right: liquid AFM image of DOTS. **c** FRAP of DOTS on the DOPC SLB surface. Left: fluorescence images showing the fluorescence recovery of the photobleached area after 120 min. Right: FRAP curve with a recovery half time t_1/2_ of 910 seconds. Scale bar=10 *μ*m **d** Schematic showing intermolecular FRET between adjacent MTS at high molecular density. **e** Schematic showing the dimension of DOTS prevents crosstalk between fluorophores, thus eliminating the intermolecular FRET. **f-g** Schematic showing the process of adding of acceptor fluorophore tagged MTS/DOTS to SLBs that were precoated with donor tagged MTS/DOTS. **h** Representative fluorescence images showing MTS acceptor (Cy3B) fluorescence signals on the SLB surface under different incubation concentrations. The MTS donor (Atto488) fluorescence intensity decreased after addition of acceptor MTS. **i** Representative fluorescence images showing DOTS acceptor fluorescence signals on the SLB surface under different incubation concentrations. The donor fluorescence intensity remained constant regardless of the density of the acceptor. **j-k** Plot showing the correlation between the fluorescence intensities of donor and acceptor MTS (**j**) or DOTS (**k**) on the SLB surface.

### The dimension of DOTS eliminates intermolecular FRET at high molecular density

We hypothesized that MTS anchored to membranes would exhibit intermolecular FRET at high molecular density, which is observed in TCR clusters. To test this concern, we anchored a conventional MTS backbone-DNA hairpin to the DOPC SLB surface. (**Fig. 1f**). Initially, a fixed quantity of Alexa 488-DNA hairpins was added to the SLB, following by washing away the excess. Subsequently, different concentrations of Cy3B hairpins were added into the SLB to tune the density. In principle, as the density of Cy3B hairpin increases, we anticipate a higher degree of intermolecular FRET, resulting in a reduction in Alexa 488 fluorescence intensity. The density of Cy3B hairpins exhibited an increase with the incubation concentration and saturated around 200 nM (**Fig. 1h** and **1j**). As expected, the fluorescence signal of Alexa 488 showed a decline as the density of Cy3B acceptor hairpins rose, providing confirmation of robust intermolecular FRET. (**Fig. 1h and 1j**). We hypothesized that our DOTS design would circumvent this problem by physically separating hairpins (**Fig. 1e**). In the 40 × 80 nm DOTS design, the nearest possible distance between fluorophores on adjacent origamis is 40 nm, which is 6-fold greater than the Forster radius and therefore effectively eliminates the possibility of intermolecular FRET occurring. We tested this claim by adding different concentrations of Cy3B-DOTS to SLBs precoated with Atto488-DOTS (**Fig. 1g**). As predicted, no fluorescence decrease in the 488 channel was observed even as the Cy3B DOTS density reached saturation at 8 nM concentration (**Fig. 1i and 1k**).

### DOTS detect TCR tension at fluid intermembrane interfaces

To measure TCR forces at fluid junctions, we proceeded by seeding naïve CD8+ T cells, isolated from OT-1 transgenic mice, onto DOPC fluid SLB coated with pMHC-loaded DOTS, in conjunction with intercellular adhesion molecule 1 (ICAM-1) (**Fig. 2a**). Upon initial contact of the T cell with the SLB, a simultaneous decrease in DOTS fluorescence intensity was observed in the cell spreading area, indicating the exclusion of DOTS from that region. Subsequently, the remaining DOTS clustered and centralized, ultimately accumulating at the center of the cell/SLB junction within a 10-minute timeframe. Concurrently, ICAM-1 tagged with GFP engaged LFA-1 and concentrated at the periphery (**Fig. 2b** and **Supplementary movie 1**). This phenomenon can be attributed, in part, to the reorganization of receptor-ligand interactions at T cell intermembrane junctions in a manner that is dependent on the height, aiming to minimize energy.^23^ However, the exclusion of origami was not solely due to axial crowding. This is supported by the observation that replacing the origami-tethered antigen with one tethered using a single DNA hairpin resulted in a significant decrease in the exclusion level, from 60% to 17%, despite the DNA hairpin being capable of extending to a greater axial height (**Fig. 2d**). We thus hypothesized the DOTS exclusion was induced by lateral crowding and was independent of protein-protein interactions. DNA origami are bulky DNA structures spanning 40 x 80 nm in the lateral dimension. The T cell expresses glycocalyx and large proteins such as CD45 phosphatase which could impose a physical constraint that sterically excludes DOTS. To confirm this, we primed T cells with anti-CD3 and then plated these cells onto blank DOTS surface lacking pMHC. Blank DOTS did not centralize due to lack of TCR engagement (**Fig. 2d, Supplementary Fig. 6**). However, they were still excluded, and the exclusion level was comparable to that of pMHC loaded DOTS. To validate the mechanism of DOTS translocation, both inward flow and outward exclusion, we performed single molecule imaging of DOTS. Here, we reduced the density of DOTS by incubating the SLB with a 1000-fold dilution of DOTS and performed timelapse fluorescence imaging during the initial stages of T cell spreading. As shown in **Supplementary movie 2**, free DOTS rapidly moved out of the cell adhesion zone, a small subset of DOTS (∼1-10) remained confined to the cell junction, and then 10’s of seconds later, TCR-ligated DOTS centralized to form what is typically described as the central supramolecular activation complex (cSMAC).

**Figure 2.**
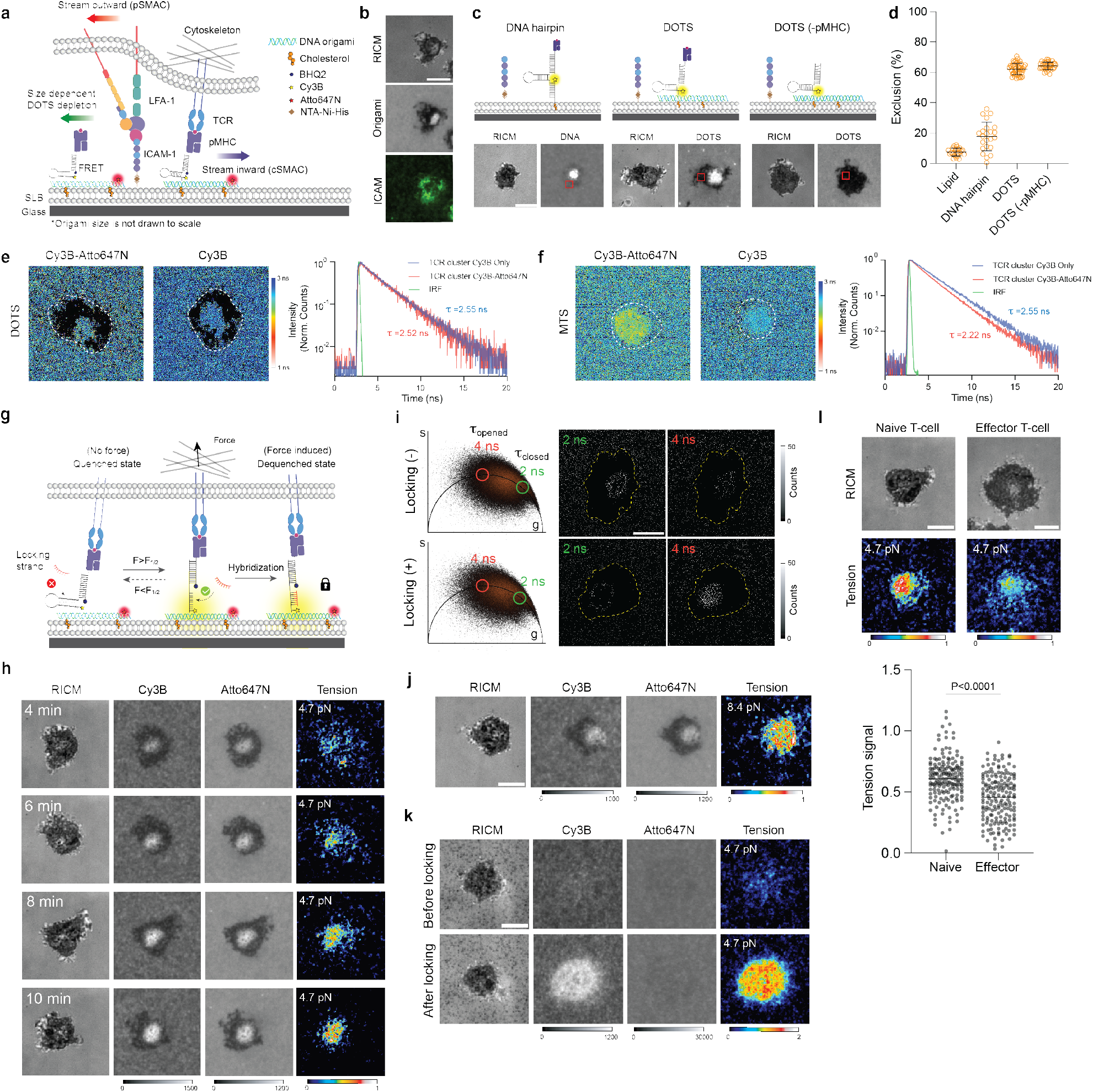
TCRs transmit mechanical force to laterally fluid antigen. **a** Schematic of functionalized SLB. Upon T cell spreading, free DOTS, TCR engaged DOTS and LFA-1-ICAM underwent reorganization at the T cell-SLB junction. **b** Representative microscope images showing the signal of T cell, DOTS and ICAM-1-GFP on the DOPC SLB surface. **c** Schematic and representative images showing different exclusions of DNA structures (hairpin, DOTS, DOTS lacking pMHC antigen) from the T cell spreading area **d** Plot comparing the exclusion levels of different DNA structures. The level of exclusion was quantified by measuring the decrease in DNA fluorescence intensity within the cell spreading area (ROIs indicated by red squares in panel (**c**)). n>23 cells for each condition. **e** Representative FLIM images of Cy3B fluorophore on the SLB surface functionalized with Atto647-DOTS and Cy3B-DOTS or Cy3B-DOTS alone. White dash lines indicate theT cell spreading areas. Black pixels in the lifetime image indicate pixels with <25 photons or lifetime > 3.2 ns (Supplementary note 1). FLIM decay curves of pixels at TCR clusters formed on Atto647/Cy3B-DOTS SLB or Cy3B-DOTS only SLB. Averaged lifetimes of pixels were noted next to the curves. **f** Representative FLIM images of Cy3B fluorophore on the SLB surface functionalized with Atto647-MTS and Cy3B-MTS or Cy3B-MTS alone. Black pixels in the lifetime image indicate pixels with <25 photons or lifetime > 3.1 ns. FLIM decay curves of pixels at TCR clusters with averaged lifetime noted. **g** Schematic showing that locking strand hybridizes to mechanically opened DNA hairpin to capture transient TCR force events. **h** Time lapse images showing the dynamics of 4.7 pN TCR tension signal in the immune synapse. **i** Comparison of *τ*_closed_ and *τ*_open_ population before and after adding locking strand. The pixel images of specific lifetime were obtained by back-projection of the points within the green circles (*τ*_closed_ population) and red circles (*τ*_closed_ population) in the phasor plots. **j** Representative RICM and tension images (8.4 pN) of T cell cultured on fluid DOPC SLB. **k** Representative images of 4.7 pN TCR tension on non-fluid DPPC SLB. **l** Representative images showing TCR tension signal of naïve and effector T cell after 20 min spreading on DOPC fluid SLB. Dot plot comparing the tension signals of naïve T cells and effector T cells. n>170 cells from 3 independent experiments. Scale bars = 5 *μ*m

After successfully establishing the ability of DOTS to interact with the TCR and lead to the formation of immune synapse, our subsequent objective was to further demonstrate DOTS’s ability to eliminate intermolecular FRET in the case of physiological TCR clusters. We seeded T cells onto SLBs coated with a mixture of Cy3B-DOTS and Atto647-DOTS and acquired the lifetime map of Cy3B fluorophore through fluorescence lifetime imaging microscopy (FLIM) (**Supplementary Fig. 7a**). FLIM measures the fluorophore excited state lifetime (*τ*), which is highly sensitive to FRET efficiency and is independent of probe density. In principle, if intermolecular FRET exists in TCR clusters, Atto647N DOTS would quench surrounding Cy3B DOTS and decrease the fluorescence lifetime of Cy3B. As anticipated, we did not observe any significant change in Cy3B fluorescence lifetime within TCR clusters compared to the background or TCR clusters formed on surfaces with Cy3B-DOTS alone, suggesting the absence of intermolecular FRET. (**Fig. 2e** and **Supplementary Fig. 7d-e**). In contrast, MTS exhibited strong intermolecular FRET as indicated by a strong Cy3B lifetime shift at TCR clusters (**Fig. 2f** and **Supplementary Fig. 7d**).

Next, we utilized DOTS to detect TCR forces by monitoring the mechanical unfolding of the DNA hairpin on the origami, which leads to dequenching of the dye (Cy3B-BHQ2) and results in a 3.3-fold fluorescence enhancement (**Supplementary Fig. 8** and **Supplementary Fig. 9a**). To deconvolve tension and cluster-induced fluorescence increases, a Atto647N fluorophore was incorporated onto the origami structure 40 nm away from the DNA hairpin to function as a density reporter as it was insensitive to force (**Supplementary Fig. 9a** and **Supplementary Fig. 3**). Tension information was obtained by quantifying the ratio between Cy3B (tension + density) and Atto647N (density). Values > 1 indicate mechanical unfolding of the hairpin and are referred to as tension signal (See **Supplementary Fig. 10** for analysis details). The force threshold of DOTS was determined by the unfolding force of DNA hairpins which depends on the GC content and stem-loop structure of the hairpin.^7^ We initially designed a DNA hairpin with 22% GC content and nine nucleotide (nt) stem to detect TCR forces exceeding 4.7 pN. TCR-pMHC interactions are highly transient with a subsecond bond lifetimes,^5^ which is difficult to capture by real time imaging (**Supplementary Fig. 9b**). In order to enhance the tension signal, we added a 15 nt ssDNA “locking strand” at a concentration of 200 nM into the cell imaging media to hybridize mechanically unfolded hairpins and lock it in the opened state (**Fig. 2g**).^24^ This locking strategy allows one to record the total accumulated mechanical events at the cell-membrane junction as long as the antigen remains bound and confined.^25^ Of note, locking strand did not cause nonspecific hairpin opening and hybridization to DNA hairpins was mechanically selective (**Supplementary Fig. 9c-e**). Within 10 min of cell-surface contact, TCR-antigen tension gradually accumulated from puncta into cSMAC (**Fig. 2h**).

We also applied FLIM to further confirm the mechanical unfolding of the DNA hairpins. Because the Cy3B/BHQ2 pair is subject to static quenching, we replaced Cy3B with Atto488 given its excellent imaging properties and larger *τ* shift after being quenched by BHQ2.^22, 26^We captured FLIM images of cells on the SLB before and after adding locking strand and generated a phasor plot for the FLIM datasets (**Fig. 2i**). From the phasor plot, we identified pixels containing fluorescence lifetimes of *τ* = 2 ns and 4 ns corresponding to folded (*τ*_closed_) and unfolded (*τ*_open_) hairpin Atto488 lifetimes (conformations), respectively,^26^ and obtained photon count maps for these two lifetimes. We then examined the change of these two populations before and after adding locking strand. As shown in **Fig. 2i**, after adding locking strand, the *τ*_closed_ population decreased at the T cell-SLB junction (indicated by yellow dash lines) whereas the *τ*_open_ population significantly increased in count. This result confirmed the accumulation of mechanically opened hairpins at the immune synapse and successfully mapped TCR-antigen mechanical events with a fluorescence lifetime readout.

Next, we adjusted the GC content of the hairpin stem to 77% to detect force with a higher magnitude of 8.4 pN. TCR forces still opened 8.4 pN DOTS and generated a tension signal comparable to that on the 4.7 pN DOTS, and thus indicating that the TCR transmits F> 8.4 pN on laterally fluid antigen (**Fig. 2j**). To further validate these findings, we substituted the DNA hairpin on the origami structure with another force-sensitive DNA element known as the tension gauge tether (TGT), which is a DNA duplex that ruptures at a specific force threshold: 12 pN in an unzipping geometry and 56 pN in a shearing geometry (**Supplementary Fig. 11a**).^27^ We observed that the cSMAC volume of T cells seeded on 12 pN TGT was smaller than that on 56 pN TGT (**Supplementary Fig. 11b**). This suggests that a subset of TCRs generated 12 pN forces, leading to the dissociation of TCR from 12 pN DOTS and subsequent translocation to the center to form the cSMAC.

To study how the fluidity of antigen influences force transmission, we replaced DOPC with DPPC lipid to create a non-fluid SLB. In this case, no clustering or exclusion of DOTS was observed as indicated by the Atto647 channel (**Figure 2k**). Notably, the tension signal on gel-phase SLBs was greater than that observed on fluid SLBs (**Fig. 2h and 2k**). This is because fluid SLBs offer little resistance to pulling in the lateral direction and confirms that TCRs experience pulling force in both shear and normal vectors when the antigen is immobilized.

We also compared the tension signal generated by naïve and effector T cells which present the same TCR but are reported to exhibit different response to antigen.^28^ Interestingly, we found the effector T cells also opened 4.7 pN DOTS but the signal intensity was much lower than that of naïve T cell (**Fig. 2l**). The differential tension signal was likely due to the difference in cytoskeletal dynamics and coupling to TCRs.^29^ We found F-actin was less dense and more heterogeneously distributed in effector T cells (**Supplementary Fig. 12**). Additionally, effector T cells exhibit a faster rate of immune synapse formation. After engaging antigen surface, TCRs on effector T cells rapidly centralized to form cSMAC within 2-3 minutes with a significant F-actin clearance (**Supplementary movie 3**). In contrast, naïve T cells took 10 min to mature the immune synapse, and F-actin clearance was less pronounced (**Supplementary Fig. 12**). The differential dynamics may also account for the discrepancy in DOTS tension signal, which reports the history of TCR mechanical events.

### Actin polymerization and membrane bending contribute to TCR forces

After establishing the presence of TCR forces at intermembrane junctions, we next employed DOTS to identify the sources of TCR forces. Prior work suggested that cytoskeleton dynamics are crucial to mechanotransduction.^30^ We treated T cells with cytoskeletal drugs to identify which component of the cytoskeleton dominates TCR force transmission at fluid interfaces. In these experiments, T cells were allowed to spread on fluid SLB surface and subjected to drug treatment for 5 min, followed by the introduction of DNA locking strand to record TCR force (**Fig. 3a**). Blebbistatin (50 *μ*M), which inhibits myosin II ATPase, did not cause a notable change in force signal compared to the vehicle control (**Fig. 3a** and **3b**). In contrast, the actin targeting drugs, CK666 (50 *μ*M) and Lat-B (5 *μ*M) dramatically decreased the force signal to 60%, and 20%, respectively (**Fig. 3b**). CK666 targets the Arp2/3 complex, thus dampening actin nucleation and branching, while latrunculin-B directly binds to actin monomers and prevents polymerization. These results suggest that myosin is less involved in TCR mechanotransduction and the cytoskeleton transmits forces to TCRs mainly through F-actin. TCRs are coupled to F-actin branches with the aid of adaptor proteins (WASP, Nck) and actin nucleating regulators (Arp 2/3) (**Fig. 3c**). We inhibited the recruitment of Nck to the TCR complex using AX-024, a small molecule that binds to the SH3 domain pocket on Nck and disturbs the interaction between Nck and CD3 subunits. When treated with AX-024, T cells exerted less tension, further confirming the role of F-actin in transmitting tension to TCR-pMHC bonds (**Fig. 3d** and **3e**).

**Figure 3.**
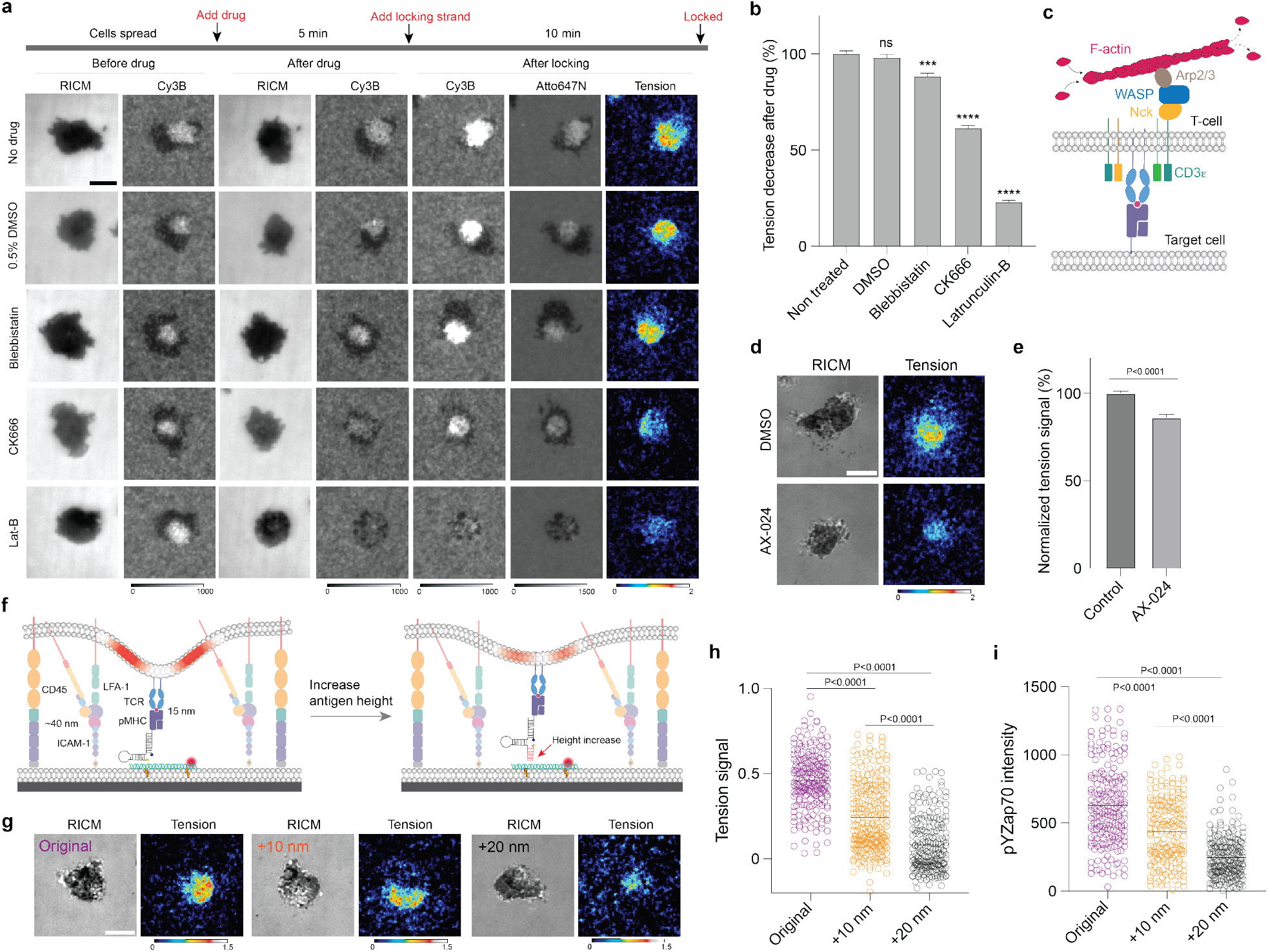
F-actin and membrane bending contribute to TCR force generation. **a** Representative microscope images showing the spreading and tension signals of T cells treated with cytoskeleton inhibitors. **b** Bar graph quantifying the tension decreases when T cell were treated with cytoskeleton drugs. n>120 cells for each group from three independent experiments. Error bar indicates SEM. ns, not significant *P*=0.4188, \*\*\**P* = 0.0001, \*\*\*\**P* < 0.0001 **c** Schematic showing the recruitment of F-actin to TCR engagement sites with the aid of adaptor proteins. **d** Representative images showing the tension signal of T cells with and without AX-024 treatment. **e** Bar graph quantifying the tension signal decrease when the T cells were treated with AX-024. **f** Schematic showing that elongating DNA hairpin increases the maximum antigen height, which decreases the membrane bending and the strain transmitted to TCR-pMHC bonds. **g** Representative images showing TCR tension signals under different antigen heights. **h-i** Dot plot quantifying intensities of TCR tension (**h**) and pYZap70 (**i**) of T cells engaging antigens with different heights. At least 200 cells from three independent experiments were analyzed. Lines indicate the mean. Scale bars = 5 *μ*m.

The mechanosensor model postulates that mechanical forces drive initial TCR triggering through a potential catch bond, which feeds into kinetic proofreading and enhances antigen discrimination. Therefore, the TCR-pMHC bond must experience force during initial TCR signaling.^5^ Paradoxically, there is minimal TCR-actin coupling prior to TCR activation. Other work also suggests that actin polymerization is not a cause but a consequence of TCR signaling^31, 32^Therefore, there must be another source of mechanical energy that is transmitted to the TCR-antigen complex to trigger initial signaling. The process of TCR-pMHC complex formation involves bringing the intermembrane space to a distance of ∼15 nm. This bond formation generates opposing forces and induces membrane bending, primarily due to the presence of large proteins at the junction. One such example is the LFA-1-ICAM-1 complex, which spans ∼36-45 nm and contributes to membrane bending at the sites of initial TCR-pMHC engagement. Furthermore, bulky proteins like the phosphatase CD45, which extends up to 40 nm in the extracellular region, may also play a role. To validate this hypothesis, we adjusted the height of the antigen by elongating the DNA hairpin on DOTS and then recorded TCR-pMHC tension change (**Fig. 3f**). All surfaces presented identical antigen and origami densities. The force threshold of the DNA hairpin was not changed by the elongated DNA tether (**Supplementary Fig.13**). T cells spread similarly across the three TCR-antigen dimensions, but the tension signal intensity significantly decreased with increasing antigen extension (**Fig. 3g** and **3h**). These differences in TCR-antigen mechanics were also associated with difference in signaling levels, as phosphorylated ZAP70 levels after 10 min of cell seeding were significantly decreased for the elongated antigen (**Fig. 3i**). Taken together, this data supports a mechanosensor model where initial forces are mediated by the size mismatch of proteins and the bending modulus of the plasma membrane at the cell-cell junction.

### Spherical SLB for investigating TCR mechanics in suspension

Although planar SLBs mimic the biochemical and biophysical properties of the plasma membrane, the surface displays a flattened topology and lacks the 3D free standing properties of target cells. In contrast, microparticles better mimic target cells and have been used as artificial antigen presenting cells to stimulate T cells and capture signaling molecules released into the immune synapse.^33, 34^Accordingly, we created a spherical SLB (SSLB) platform by coating a 5 *μ*m particle with SLB and then tethered DOTS onto it to measure TCR forces at T cell/SSLB junctions (**Fig. 4a**). In a typical experiment, DOTS-SSLBs and T cells were mixed at 1:1 ratio in suspension in the presence locking strand and imaged on a confocal microscope after 30 min incubation. TCR engaged DOTS and translocated to the center to form cSMAC (**Fig. 4b**). Similar to the approach used with planar SLB DOTS, we extracted tension information by analyzing the Cy3B/Atto647N ratios (**Supplementary Fig.14**). At the T cell-SLB junctions, the average Cy3B/Atto647N ratio was found to be 2.2-fold higher and returned to background levels when the locking strand was absent. This indicated the presence of 4.7 pN forces that caused the opening of hairpins at the junction (**Fig. 4c**). We next conducted z-scanning across the junction and performed image arithmetic on successive focal planes to construct the 3D view of tension signal. As shown in **Fig. 4d** and **Supplementary movie 4**, the tension signal covered the whole synapse and overlapped with the DOTS central clusters. Taken together, T cells generated and transmitted F >4.7 pN to TCR-pMHC complexes, even if the pMHC was laterally fluid and tethered to a microparticle in suspension.

**Figure 4.**
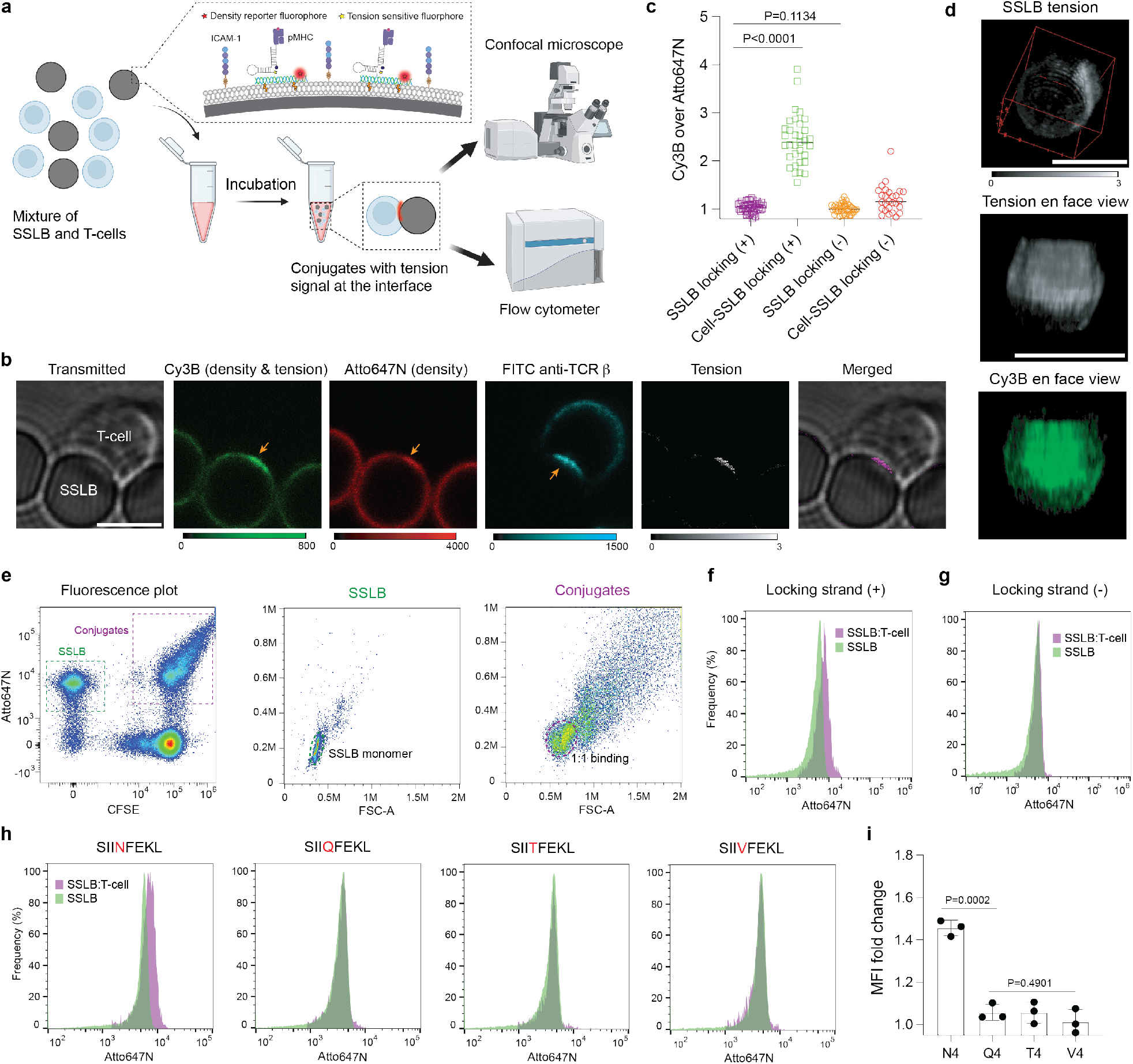
SSLB as a platform to study 3D TCR tension. **a** Schematic showing the workflow of using SSLBs to study TCR mechanics through high resolution confocal microscopy and high throughput flow cytometry. **b** Representative images showing the spatial distribution of DOTS, TCR, and tension signal at the middle layer of the T cell-SSLB junction. **c** Plot comparing the Cy3B/Atto647N ratios on SSLB surfaces and SSLB-T cell junctions with and without locking strand. Data from each group was acquired from >25 cells **d** 3D reconstructed image showing the 3D distribution of DOTS and tension signals at the junction. **e** Flow cytometry fluorescence dot plot (Atto647 vs CFSE) showing T cells, SSLBs (green dash line), and conjugates events (purple dash line). Forward vs side scatter plot was used to gate out SSLB monomers (green dash line) and T cell-SSLB conjugates at 1:1 binding stoichiometry (purple dash line). **f** Atto647N fluorescence histograms of SSLBs (green) and 1:1 T cell/SSLB conjugate (purple) events. **g** Atto647N fluorescence histograms of SSLBs and 1:1 T cell-SSLB conjugates when locking strand was absent. **h** Atto647N fluorescence histogram of SSLBs and 1:1 T cell/SSLB conjugates where SSLBs were modified DOTS presenting antigens with different potencies. **i** Plot quantifying the Atto647N mean fluorescence intensity (MFI) difference between 1:1 T cell/SSLB conjugates and SSLBs. \*\*\**P* < 0.001 \*\*\*\**P* < 0.0001. Scale bars = 5 *μ*m.

One advantage of the DOTS-SSLB platform is that each particle is at the cellular scale and thus amenable to flow cytometry analysis and enables high throughput quantification of TCR forces. In suspension, three distinct populations were present-single T cells, single SSLBs, as well as conjugates of the two entities with various stoichiometries. The SSLBs were identified with the DOTS fluorescence signal (Atto647N). Note that here we used the Atto647N/BHQ2 instead of Cy3B/BHQ2 dye pair to decorate hairpin because Atto647N is better suited for flow analysis and displays minimal spectral overlap with CFSE dye used to label the T cells (**Fig. 4e**). Events that showed dual positive signals (Atto647N+CFSE) were identified as SSLB-T cell conjugates (purple box in **Fig. 4e**), from which 1:1 stoichiometry conjugates were selected using the forward scattering profiles for comparison with single SSLBs. In principle, TCR forces would light up the DOTS on the SSLB surface, making conjugate events exhibit a stronger Atto647N signal than individual SSLB events. Of note, unlike microscopy, flow cytometry measures the integrated fluorescence intensity of particles, thereby obviating the need for ratiometric analysis. We analyzed 200,000 events and observed a 40% mean fluorescence intensity (MFI) difference between these two populations (**Fig. 4f**). This MFI change was due to tension as control experiments without locking strand did not show notable difference (**Fig. 4g**).

The high throughput readout of SSLB signal enables the use of this platform as a potential tool for rapid antigen screening. It has been demonstrated that mechanical forces are highly correlated with antigen potency and subsequent T cell activation, and in fact the force level is likely a better measure of antigen potency compared to affinity.^35, 36^ To demonstrate this concept, we mutated the fourth amino acid of the cognate OVA derived peptide SIINFEKL (N4) to obtain SIIQFEKL (Q4), SIITFEKL (T4), SIIVFEKL (V4) altered peptide ligands (APLs). DOTS were modified with APLs and tethered to SSLBs to engage T cells. Compared to cognate N4 antigen, APLs SSLB displayed a lower conjugation efficiency to T cells (**Supplementary Fig. 15**) and did not show any fluorescence change after engaging T cells (**Fig. 4h-i**). This shows that the TCR-antigen tension signal at the intermembrane junction is highly specific to the agonist antigen and confirms the potential of flow analysis of DOTS to be used as a tool for antigen screening.

### DOTS revealed TCR tension at the physiological T cell-B cell junctions

There is always the question of whether the native TCR-pMHC complex experiences the same magnitude of pN forces as those recorded on synthetic surfaces. Membrane protrusions, contractility of target cells, and proteins on the target cell membrane could alter the forces experienced by the TCR-pMHC complex. Thus, we next aimed to anchor DOTS onto target cell membranes to study forces at authentic cell-cell junctions. We screened different conjugation chemistries including maleimide-thiol, streptavidin-biotin, and cholesterol to anchor DOTS to cell membranes (**Supplementary Fig. 16 a-c**).^37, 38, 39^ Origami labeling efficiency was dependent on the number and locations of the binding sites.^40^ Overhangs near the edge of the origami sheet offered more robust tethering compared to overhangs in the middle. Therefore, we introduced 12 additional anchor strands at the edges of DOTS (**Supplementary Fig.1**). In addition, because glycocalyx hindered origami access to the membrane, ^41^ we used a long “bridge strand” to link the cholesterol strand and anchor strand on the DOTS to help minimize steric crowding at the plasma membrane (**Fig. 5a**). We labeled three target cell lines (EL-4, B16-F10 and B cells) and ended up selecting resting murine B cells for the following studies because they have a comparable size to T cells, exhibit high binding efficiency and simple binding stoichiometry with T cells (**Supplementary Fig. 16d-f**). With all these efforts, DOTS anchored to B cell membrane homogenously distributed across the surface and maintained its integrity for hours at RT (**Supplementary Fig. 17**).

**Figure 5.**
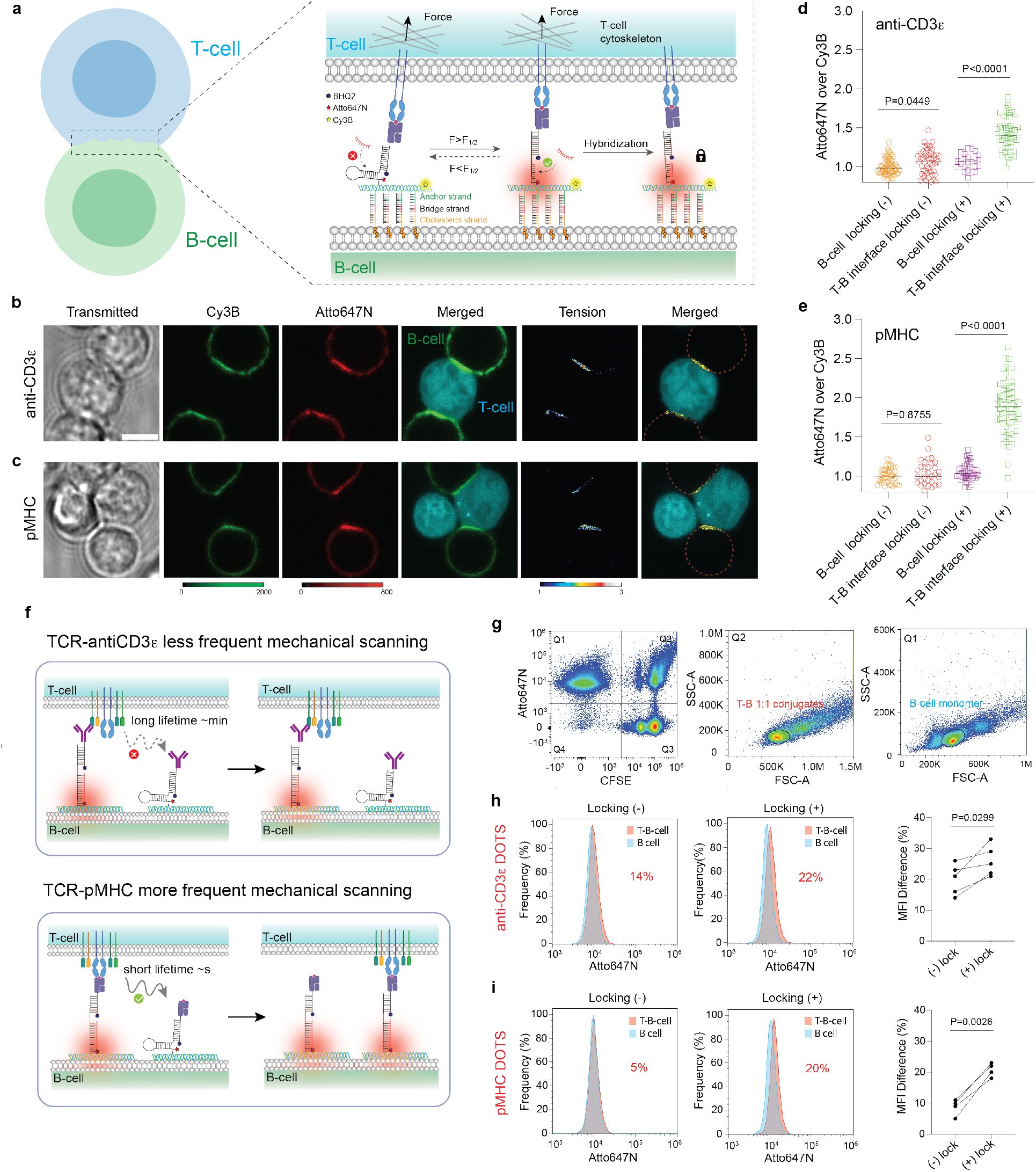
TCR tension at T-B cell interfaces. **a** Schematic showing the functionalization of B-cell membrane with DOTS. Locking strand was used to capture the tension signal at the T-B cell interface. **b-c** Representative images showing the DOTS and tension signals at the T-B cell interfaces. DOTS were modified with anti-CD3*ε* (**b**) or pMHC (**c**). **d-e** Plot comparing the Atto647N/Cy3B ratios on the B-cell surface and T-B interfaces with and without locking strand after 30 min incubation. B-cells are modified with anti-CD3*ε* DOTS (**d**) or pMHC DOTS (**e**). Data from each group was acquired >25 T-B cells. **f** Schematic showing that TCR-anti-CD3*ε* has a higher bond lifetime than TCR-pMHC and thus scans and lights up fewer DOTS within a specific period (30min). **g** Flow cytometry fluorescence plots and forward scatter plots were used to gate out the B-cell monomer and T-B 1:1 binding population. **h-i** Atto647N fluorescence histograms of B-cells (blue) and T-B 1:1 conjugates (red) with and without locking strand. Plot quantifying Atto647N MFI difference between B-cells and T-B cell 1:1 conjugates with and without locking strand present in the media. Data were acquired from 4 independent experiments and 4 independent mice. B-cells are modified with anti-CD3*ε* DOTS (**h**) or pMHC DOTS (**i**). Scale bars = 5 *μ*m.

To measure TCR forces, we loaded anti-CD3*ε* on 4.7 pN DOTS before anchoring to the B-cell membrane. DOTS modified B cells were mixed with T cells and allowed to engage for 30 min before imaging and analysis. We found that B cells strongly interacted with T cells and accumulated DOTS at the interface (**Fig. 5b**). TCR forces separated Atto647N from the quencher, leading to a fluorescence increase over the force insensitive label of Cy3B. We analyzed 25 T-B conjugates and obtained an average of 5 ± 16% Atto647/Cy3B ratio increase at the T-B cell junctions compared to that on B-cell surfaces (**Fig. 5d**). Note that this change corresponded to real-time TCR forces as no locking strand was added to lock the mechanically unfolded hairpin. After introduction of locking strand (200 nM), the ratio further increased at the T-B cell junctions by 43 ± 20% (**Fig. 5c** and **5d**). These results demonstrate that the TCR transmits >4.7 pN forces to its antiCD3*ε* ligands at cell-cell junctions. We next labeled B-cells with DOTS presenting pMHC antigen to visualize the force at TCR-pMHC bonds. In contrast to anti-CD3e, in the absence of locking strand, we did not observe a statistically significant Atto647N/Cy3B ratio change at the T-B cell junction. However, Atto647/Cy3B ratio increased by 90 ± 35% when the locking strand was added (**Fig. 5c** and **5e**) and even surpassed that of the antiCD3*ε* DOTS signal. In retrospect, this finding is consistent with the known short lifetime but high frequency sampling and scanning that is inherent to TCR-pMHC interactions which leads to greater accumulated signal at the T-B cell interface (**Fig. 5f**).

We next used flow cytometry to analyze the tension signal in high throughput following the methods described under the SSLB-T cell measurements in **Fig. 4**. 1:1 T-B conjugate and B-cell monomer populations were gated out from the fluorescence and forward scatter plots, and their Atto647N intensities were compared. In the absence of locking strand, we observed a 20 ± 5% MFI change in anti-CD3*ε* DOTS coated B-cells after its binding to T cells (**Fig. 5h**). Consistent with the microscopy data, a smaller change of 9 ± 3% was observed for pMHC-DOTS coated B-cells (**Fig. 5i**). However, in the presence of locking strand, the MFI change significantly increased from 9% to 21 ± 2% for pMHC-DOTS modified B-cell (**Fig. 5i**). An increase was also observed in antiCD3*ε* but was less pronounced than that of pMHC-DOTS (**Fig. 5h**). Collectively, the findings from both microscopy and flow cytometry analyses demonstrated the presence of >4.7 pN mechanical events at immune cell-cell junctions, with their characteristics modulated by the affinity of TCR ligands.

## Discussion

In this work, we developed DOTS by combining the idea of molecular tension sensors and DNA origami nanodevices and established its performance in investigating TCR mechanotransduction at intermembrane junctions. This contrasts with previously reported tension sensor designs which suffered from intermolecular FRET and displayed many supraphysiological characteristics regarding stiffness and either ultrahigh or limited lateral diffusion. With DOTS, we demonstrated that TCRs transmit 5-10 pN forces to its antigen at fluid interfaces. Forces were not only characterized by widely used widefield microscope, but further validated using fluorescence lifetime-based imaging as well as flow cytometry with a high throughput readout.

It has been long assumed that the endogenous force transmitted to receptor-ligand bonds are generated by the cell cytoskeleton. In this report, we identified another source of force caused by the protein size mismatch at the immune synapse. TCR-pMHC complexes, displaying a shorter size compared to CD45 and LFA-1-ICAM-1, must “pinch” the intermembrane junction bringing it closer in order to interact, which causes strain in the interaction to trigger TCR signaling. The well-documented cell cytoskeleton does contribute to TCR forces but only after initial triggering and proximal kinase activation. Moreover, cytoskeletal force generation and transmission to the TCR is mediated by F-actin dynamics and is less reliant on myosin activity. This finding mirrors the published work that F-actin maintains synapse persistence whereas myosin is dispensable in synapse formation.^42, 43^

It is worth noting that DOTS are highly programmable. For example, DOTS were tethered to microparticles creating a spherical tension sensor platform that demonstrated a correlation between force intensity and antigen potency and provided a new approach for antigen screening. Additionally, we successfully functionalized target cell membrane with a homogenous DOTS layer which allowed us to investigate the TCR tension at T cell-target cell junctions. Previously, DNA hairpin structures were inserted onto epithelial cells to study E-cadherin mediated tensile force.^39^ However, along with intermolecular FRET issue, this design is susceptible to endocytosis as DNA hairpins are only composed of three short DNA strands. In contrast, DOTS preserved their structural integrity for hours and no internalization was observed. Moreover, ideally one can functionalize the DNA origami with different ligands through straightforward DNA hybridization to investigate the mechanical communication between receptors. Prior work reported that CD28 engagement increases traction forces associated with CD3.^44^ To explain this finding at the molecular level and validate the programmability of DOTS in force sensing, we labeled anti-CD28 with DNA and incorporated it to the DOTS platform to investigate how CD28 engagement tunes the forces experienced by individual TCR-antigen bonds. Consistently, we observed a stronger TCR tension signal when CD28 was ligated (**Supplementary Fig.18**). One limitation of DOTS pertains to its low signal to background ratio in detecting real-time TCR tension, but this limitation will likely be resolved by introducing additional FRET pairs onto the DOTS so that TCR tension would lead to separation of multiple FRET pairs with a stronger fluorescence response.

## Materials and Methods

### Reagents

DOPC (Cat# 850375C-200mg), Ni-NTA-DGS (Cat # 790404C-5 mg), DPPC (Cat# 850355C-25mg), 18:1 Biotinyl Cap PE (Cat# 870273C-25mg), were purchased from Avanti Polar Lipids Inc. (Alabaster, AL). Heat-inactivated fetal bovine serum (FBS) (Cat# 35-015-CV), penicillin-streptomycin solution (Cat# 30-234-CI), and gentamicin sulfate solution (Cat# 30-005-CR) were purchased from Corning Mediatech (Corning, NY). 1M Tris (Cat#: AM9856), 0.5 M EDTA (Ca#: AM9260G), 1M MgCl_2_ (Cat# AM9530G), RPMI (Cat#11835030), Texas Red™ DHPE (Cat# T1395MP), human IL-2 (Cat# PHC0026) and CellTrace™ CFSE (Cat# C34570) were purchased from ThermoFisher (Waltham, MA). Bovine serum albumin (BSA) (Cat# 10735078001), Latrunculin B (Cat# L5288, >80%), Atto647N NHS ester (Cat# 18373-1MG-F), Atto488 NHS ester (Cat# 41698-1MG-F), 100 kDa Amicon ultra-0.5 centrifugal filter (Cat# UFC510096) and Hank’s balanced salt solution (H8264-6X500ML) were purchased from Sigma Aldrich (St. Louis, MO). Cy3B NHS ester (Cat# PA63101) was purchased from GE Healthcare (Pittsburgh, PA). CK666 (Cat# ab141231) was purchased from Abcam (Cambridge, United Kingdom). Blebbistatin (Cat# 72402) was purchased from STEMCELL (Vancouver, Canada). Streptavidin (Cat# S000-01) was purchased from Rockland Immunochemicals Inc. (Rockland, NY). Biotinylated pMHC ovalbumin (SIINFEKL) was obtained from the NIH Tetramer Core Facility at Emory University. P2 size exclusion gel (Cat#1504118) was purchased from Bio-Rad (Hercules, CA). 3 mL syringes were purchased from BD bioscience (San Jose, CA). Cell strainers (Cat# 15-1100) were bought from Biologix (Shandong, China). Midi MACS (LS) startup kit (Cat# 130-042-301) (separator, columns, stand), mouse CD8+ T cell isolation kit (Cat# 130-104-075), and resting mouse B-cell isolation kit (Cat# 130-090-862) were purchased from Miltenyi Biotec (Bergisch Gladbach, Germany). Oligonucleotides were obtained from Integrative DNA Technologies (Coralville, IA) and Biosearch Technologies (Hoddesdon, United Kingdom). His-ICAM-1 (Cat# 50440-M03H) was purchased from Sino Biological (Beijing, China). Anti-mouse CD28 (Cat# 102102), biotinylated anti-mouse CD3e (Cat# 100304) and FITC anti-mouse TCRb antibody (Cat# 109206) were purchased from BioLegend (San Diego, CA). Azide-PEG4-NHS ester (Cat# AZ103-100) and Sulfo-DBCO NHS ester (Cat# A124-10) were purchased from Click Chemistry Tools (Scottsdale, AZ). Alexa 488 Mouse Anti-ZAP70 (pY319)/Syk (Y352) (Cat# 557818) was purchased from BD Biosciences (Franklin Lakes, NJ). Single-stranded scaffold DNA, type p7560 was purchased from tilibit nanosystems (Munich, Germany).

### Harvest and purification of primary naïve OT1 T cell

OT-1 T cell receptor transgenic mice were bred and housed at Emory University’s Division of Animal Resources Facility, following the guidelines of the Institutional Animal Care and Use Committee. The OT-1 T cells express the CD8 co-receptor and have a specific recognition for the chicken ovalbumin epitope 257–264 (SIINFEKL) in the context of the MHC allele H-2K. Naïve OT-1 T cells were isolated from the spleen using magnetic activated cell sorting, as instructed by the manufacturer’s CD8+ T cell Isolation Kit (Miltenyi Biotec, Germany). In brief, a single cell suspension of splenocytes was obtained and incubated with biotinylated antibodies targeting unwanted splenic cell populations. These populations were separated from the OT-1 T cells using anti-biotin magnetic beads and enrichment on a magnetic column. The purified T cells were then washed, suspended in HBSS solution, and kept on ice until the experiment.

### Preparation of effector OT1 T cells

Splenocytes from OT-1 transgenic mice were pulsed with 100 nM OVA peptide in RPMI media containing 10% FBS for 2 days. Afterwards, activated lymphoblasts were purified by density gradient centrifugation and adjusted to a concentration of 1 million cells/mL in RPMI media containing 10% and 30 IU/mL IL-2. Cells were then maintained and split as needed in RPMI media containing 10% FBS and 30 IU/mL IL-2 until imaging on day 7.

### Retroviral transduction

To generate retrovirus, Phoenix E cells were transfected with expression vectors (Life-act) and packaging plasmids (kindly provided by Morgan Huse lab at Sloan Kettering Institute) using the calcium phosphate method. After 48 hours at 37°C, viral supernatants were collected and added to OT1 blasts two days following peptide stimulation. The mixtures were centrifuged at 1400 × g in the presence of polybrene (4 µg/ml) for 2 hours at 35°C. T cells were then split at a 1:3 ratio in medium containing IL-2 and cultured at 37°C. After overnight culture, selection was conducted to removed untransduced T cells. The remaining T cells were further cultured for additional two days before use.

### Synthesis of dye-labeled DNA strands

Oligonucleotide-dye conjugates were prepared by coupling the amine on the DNA strand with activated NHS-ester of the organic dye. Briefly, aminated DNA strands (100 µM) was mixed with excess Cy3B-NHS ester, Atto647-NHS ester or Atto488-NHS ester (500 µg/mL) and allowed to react in aqueous solution (pH=9) for 3 hours at room temperature. The mixture was then filtered by P2 gel to remove salts and unreacted dyes and then purified by HPLC with an Agilent AdvanceBio Oligonucleotide C18 column (4.6 x 150 mm, 2.7 µm). The mobile phase A: 0.1 M TEAA and B: ACN were used for a linear gradient elution of 10-100% B over 50 min at a flow rate of 0.5 mL/min. The desired products were characterized by ESI mass spectrometry (**Supplementary Fig1** and **Fig.2**).

### Preparation of DOTS

DOTS was assembled by mixing p7560 DNA scaffold strand (30 nM), eight anchor strands (300 nM), dye modified DNA hairpin strand (600 nM), dye modified density reporter strand (600 nM), ligand strand (1500 nM) and other 82 staple strands (300 nM) in the folding buffer (5 mM Tris, 1 mM EDTA, 8 mM MgCl_2_). Anchor strands were elongated at its 5’ end with 32 bases complementary to the DNA scaffold. DNA hairpin strand was elongated at its 3’ end with 32 bases complementary to the DNA scaffold. Note that DOTS for cell-cell experiments had additional 12 anchor strands. Sequences of these strands were shown in **Table S2-S4**. The locations of these strands on the origami platform were shown in **Supplementary Fig. 3**. DNA origami were annealed by heating at 90 °C for 15 min and cooling down to 4 °C at a rate of 1 °C/min. Afterwards, excess oligonucleotides were removed via ultrafiltration (100 kDa 0.5 mL Amicon Ultrafilters). Purified origami structures were stored at -30 °C in PBS supplemented with 8 mM MgCl_2_ and used within one week. Successful assembly of DNA origami was confirmed through agarose gel electrophoresis (0.75 % agarose gel, 0.5 X TBE buffer, 8 mM MgCl_2_). Agarose gel was run on ice for 2 hours at 70V and stained with ethidium bromide.

### Preparation of small unilamellar vesicles (SUVs)

Lipids with desired composition were mixed in a round-bottom flask. The lipid mixture was dried using a rotary evaporator to remove the chloroform. The lipids were further dried under a steam of compressed N_2_ and then hydrated with PBS to a concentration of 2 mg/mL. Three cycles of freeze-thaw were performed to disrupt large, multilamellar vesicle suspensions. The resulting lipids were then repeatedly extruded through an 80-nm polycarbonate membrane filter until the solution became clear (∼10 times) and stored at 4°C before use.

### Preparation of functionalized planar SLB

The wells in optically transparent 96-well plates (ThermoFisher) were washed with 5 mL ethanol and water and etched with 6.5 M NaOH for 1 hour at room temperature. The etched wells were washed with 10 mL water and treated with 100 *μ*L 0.5 mg/mL SUVs for 5 min. SUVs containing 98% DOPC and 2% DGS-NTA (Ni) lipids were used for making fluid phase SLB, and SUVs containing 98% DPPC and 2% DGS-NTA (Ni) lipids were used for non-fluid phase SLB. After treatment, unbounded vesicles were removed by washing with 5 mL PBS. SLBs were subsequently blocked with bovine serum albumin (BSA, 0.05%) in PBS for 30 min and washed with 5 mL PBS. Then, cholesterol DNA strands (250 nM) was added to SLB, incubated for 1 hour and rinsed with PBS. Subsequently, DOTS (5 nM) was added for 1 hour to bind to cholesterol strands on the SLB. The wells were then washed with PBS supplemented with 8 mM MgCl_2_ to remove excess DOTS. Streptavidin (5 *μ*g/mL) and pMHC ligands (5 *μ*g/mL) were added sequentially to the SLB and incubated for 45 min followed by washing with PBS supplemented with 8 mM MgCl_2_. Finally, His-tagged ICAM-1 (1 *μ*g/mL) was added for 1 hour (resulting in a molecular density of 100 molecules/*μ*m^2^). Wells were then washed with PBS supplemented with 8 mM MgCl_2_ and buffer-exchanged with Hank’s balanced salt solution (HBSS) before adding cells.

### Preparation of DOTS functionalized spherical SLB

Transfer 10 *μ*L non-functionalized silica beads (5.00 µm diameter, 10% w/v) into a 1.5 mL microcentrifuge tube and wash with PBS twice with bench centrifuge. SSLBs were formed by incubating beads with 500 uL 0.5 mg/mL SUVs (2% DOGS-NTA and 98% DOPC) on the rocker for 30 min at room temperature. The resultant SSLB were washed three times with PBS by centrifuging at 300g for 3 min and then blocked with 0.05% BSA for 30 min. After three PBS washes, the SSLB were resuspended into 250 nM cholesterol strand, incubated for 1 hour on the rocker at room temperature and washed with PBS. Meanwhile, to DOTS solution, 20-fold streptavidin was added for 40 min to functionalize DOTS with streptavidin. Streptavidin modified DOTS was purified via ultrafiltration (100 kDa 0.5 mL Amicon Ultrafilters) and incubated with 20-fold biotinylated pMHC for 40 min. Without further purification, pMHC functionalized DOTS were added to above cholesterol strand coated SSLBs at a concentration of 5 nM and incubated for 1 hour at room temperature. The DOTS coated SSLBs were then washed with PBS supplemented with 8 mM MgCl_2_ to remove excessive DOTS. Finally, DOTS-SSLB were incubated with His-tagged ICAM-1 (1 *μ*g/mL) for 1 hour, washed with PBS supplemented with 8 mM MgCl_2_ and rinsed with HBSS before being mixed with T cells. A total of 300,000 DOTS-SSLB particles were combined with 300,000 T cells in a 300 *μ*L HBSS, with the mixing taking place either in a 96-well plate for confocal microscopy characterization or in an Eppendorf microcentrifuge tube for flow cytometry characterization. To capture the tension signal, the locking strand with a concentration of 200 nM was introduced into the medium. The mixture was incubated at RT for 30 minutes before proceeding with imaging and flow analysis.

### Quantitative fluorescence microscopy

Surface density of DOTS and ICAM-1 was measured using a quantitative fluorescence microscopy technique developed by Groves and others.^45^ Briefly, SUVs containing 0.1 mole percent (mol %) Texas Red-DHPE (TR-DHPE) and 99.9 mol % DOPC were mixed to generate vesicle mixtures with TR-DHPE ranged from 0 to 0.1 mol %. These solutions were diluted to 0.5 mg/ml and added to a cleaned 96-well plate to establish a lipid calibration curve. The lipid head has an area of ∼0.72 nm^2^, allowing for ∼2.78 × 10^6^ lipid molecules to be packed in 1 *μ*m^2^ surface. Given the ratio of the TR-DHPE in the lipid vesicle, the TR-DHPE density could be calculated. We measured the fluorescence intensities of the SLB with different ratios of TR-DHPE and generated a calibration curve based on this (**Supplementary Fig. 5a**). To determine the density of samples (e.g. DOTS or ICAM-1 ligand), a scaling factor (F factor) was introduced to account for the difference in brightness between the sample fluorophores and TR-DHPE. We prepared varying concentrations (50 to 200 nM) of TR-DHPE liposome and samples and plotted the fluorescent intensity against concentration (**Supplementary Fig. 5b**). The slope of sample was directly compared to the TR-DHPE to yield a F factor. The F factor was subsequently used to infer the molecular density of the sample from the SLB calibration curve.

### Labeling B-cells with DOTS

Resting B-cells were enriched from the spleen using the B-cell Isolation Kit (Miltenyi Biotec, Germany, Cat# 130-090-862). 1 mL 8 million/mL B-cells in HBSS buffer were sequentially incubated with 1 *μ*M cholesterol strand (30 min, room temperature), 1 *μ*M bridge strand (30 min, 4 °C), 1 *μ*M fortifier strand (30 min, 4 °C) and washed with PBS after each incubation. In a separate tube, DOTS were functionalized with streptavidin and biotinylated ligand (anti-CD3*ε* or pMHC) sequentially at a 20-molar excess for 45 min at room temperature. DOTS were purified after each functionalization via ultrafiltration (100 kDa 0.5 mL Amicon Ultrafilters). Finally, ligand functionalized DOTS were added to DNA strand modified B-cells at a concentration of 35 nM in PBS (supplemented with 0.5 % BSA and 8 mM MgCl_2_) and incubated for 1 hour at 4 °C followed by two washes with HBSS to get DOTS modified B cells. In an Eppendorf microcentrifuge tube, 300,000 DOTS-B cells were combined with 300,000 CFSE-stained T cells in 300 *μ*L HBSS. The mixture was gently centrifuged at 94 rcf for 5 minutes and incubated for 10 minutes at RT, allowing the T cells and B cells to come into contact and form conjugates. Afterwards, the mixture was gently pipetted to disrupt any nonspecific conjugates. Subsequently, the locking strand with a concentration of 200 nM was introduced into the medium. The mixture was incubated at RT for 30 minutes before proceeding with imaging and flow analysis.

### Immunostaining

A total of ∼1 × 10^5^ T cells cultured on 96 well plate surfaces were fixed by 4% formaldehyde in PBS for 10 min. The surfaces were gently washed with PBS to remove the formaldehye. Cells were then permeabilized in 0.1% Triton X-100 for 5 min and washed with PBS. Subsequently, 2% BSA was added to the surfaces and incubated overnight at 4°C. On the next day, the surfaces were washed with PBS. 20 uL Alexa 488 Mouse Anti-ZAP70 (PY319)/Syk (PY352) (Cat# 557818 from BD Biosciences) was mix with 80 uL staining buffer (0.5% BSA in PBS) and added to the surface for 1 hour at room temperature. Surfaces were then washed with PBS before imaging on total internal reflection fluorescence (TIRF) microscope. For actin staining, after fixation and permeabilization, the cells were incubated with 1X phalloidin-ifluor488 (Cat# ab176753 from Abcam) in 1% BSA for 1 hour at room temperature and washed with PBS before imaging on TIRF.

### DNA-anti-CD28 conjugation via click chemistry

300 uL 0.5 mg/mL anti-mouse CD28 was loaded into Amicon ultrafilters (0.5 mL, 100 KDa) and spun at 14, 000 g for 5 min at 4 °C. The volume was adjusted to 100 μL with PBS, resulting in an approximate concentration of 1.5 mg/mL. Next, Sulfo-DBCO-NHS reagent was resuspended into DMF at a concentration of 30 mM. The reagent was then added to the antibody solution at a molar ratio of 10:1 and incubated on ice for 2 h. Excessive Sulfo-DBCO-NHS was then removed using 7K MWCO Zeba Spin Desalting columns (ThermoFisher). Azide modified DNA strands was added to DBCO modified anti-CD28 at a molar ratio of 12:1. The reaction was incubated overnight at 4 °C. On the following day, the DNA-anti-CD28 mixes were added to Amicon ultrafilters (0.5 mL, 30 KDa) and washed 8 times to remove unreacted azide DNA strands. Conjugation was confirmed using SDS-PAGE gel (**Supplementary Fig.18**). The concentration of DNA-antiCD28 was determined using Micro BCA protein Assay kit (ThermoFisher).

### Atomic force microscopy imaging

After purification, DNA origami structure was confirmed through AFM imaging. Imaging was conducted on Bruker MultiMode NanoScope V AFM using tapping mode in liquid with a Bruker ScanAsyst-Fluid+ cantilever. For sample preparation, DNA origami was first diluted to 1 nM in TE-Mg buffer (5 mM Tris,1mM EDTA,10 mM MgCl_2_) and then 20 uL of diluted origami was added onto freshly cleaved mica precoated with 0.1 ug/ml poly-L-ornithine.

### Fluorescence imaging

Epi fluorescence microscopy and TIRF microscopy experiments were performed on a Nikon Eclipse Ti inverted microscope driven by the NIS Elements software. The microscope features an Evolve electron-multiplying charge-coupled device (Photometrics), an Intensilight epifluorescence source (Nikon), a CFI Apo 100 X (numerical aperture 1.49) objective (Nikon) and a total internal reflection fluorescence launcher with three laser lines: 488 (10 mW), 561 (50 mW), and 638 nm (20 mW). This microscope also includes the Nikon Perfect Focus System, an interferometry-based focus lock which allows the capture of multi-point and time-lapse images without loss of focus. Cell-SSLB and cell-cell experiments were imaged on a Nikon confocal microscope. This microscope is equipped with a 60 X oil objective and a C2si scan head. Experiments were performed using three laser lines (488 nm, 561 nm and 640 nm) and the filters with the following bandpasses: 445/50+60LP, 525/50, and 600/50 nm. Z-stack imaging was performed using the ND Acquisition module in Nikon Elements.

### Fluorescence lifetime imaging microscopy (FLIM)

FLIM imaging for intermolecular FRET experiments (**Fig. 2e-f**) was performed on a Nikon Ti Eclipse Inverted confocal microscope with a Plan Apo Lambda 60?/1.40 Oil objective. The confocal microscope is equipped with a Picoquant Laser Scanning microscope TCSPC Upgrade with SymPhoTime 64. Samples were excited with a 40 MHz pulsed 485 ± 10 nm laser. The laser light was reflected using a 560 nm dichroic filter and the detector collected emitted photons that passed a 582/75 nm bandpass filter. Tension FLIM (**Fig. 2l**) was performed on Timebow imaging on Abberior STED microscope equipped with 60?/1.40 Oil objective, two pulsed STED lasers (595 and 775 nm), four excitation lasers (405, 485, 561, 640 nm) and a MATRIX array detector.

### Flow cytometry analysis

Flow cytometry experiments were conducted on CytoFlex flow cytometer (Beckman coulter) which features 488 nm and 638nm laser lines and bandpass filters (450/45, 585/42, 660/10 nm).

### Statistical analysis

All experiments were conducted as at least three technical and biological replicates. Statistical significance was determined on GraphPad Prism using either one-way analysis of variance or two-tailed student’s *t* tests. Data were presented as bars with mean±SEM or scatter plots with line representing mean.

## Acknowledgements

K.S. acknowledges the financial support from NIH R01 AI172452 and R01 GM131099. Y.H. is a recipient of the National Cancer Institute Predoctoral to Postdoctoral Fellow Transition Award (F99CA274690). Y.D. is a recipient of American Heart Association Postdoctoral Fellowship (23POST1028975). We thank the National Institutes of Health (NIH) Tetramer Facility at Emory University for providing the biotinylated pMHC monomers. We thank Dr. David Dunlap for the help with AFM imaging. We thank Joseph Mancuso and Dr. Hiroaki Ogasawara for helping run ESI-Mass Spec on dye conjugated oligos. This research project was supported in part by the Emory University Integrated Cellular Imaging Core. The content is solely the responsibility of the authors and does not necessarily reflect the official views of the National Institute of Health.

## Author contributions

Y.H. and K.S designed research. Y.H. performed experiments and analyzed the data. Y.D designed DNA origami structure and helped assemble and purify DNA origami. A.V. conducted computational modeling experiments to analyze the length and mechanical properties of DOTS. S.N. helped conduct FLIM imaging and analyzed the FLIM data. J. R. helped with T-cell purification. Y.H. and K.S. wrote the manuscript, with all the authors providing inputs.

## Notes

### Competing Interest Statement

The authors have declared no competing interest.

### Summary of Updates

Updated supporting information and supplementary videos

