## Supporting information for "DNA Origami Tension Sensors (DOTS) to study T cell receptor mechanics at membrane junctions"

#### Table of content

**Figure S1** Synthesis and purification of dye-labeled targets……………………………………..……..3

**Figure S2** Mass spectrometry characterization of dye-labeled oligo targets..……………...……….…4

**Figure S3** Schematics of DNA origami structures…………...……………………………………………..7

**Figure S4** Stability analysis of DNA origami under different cell imaging conditions……………..……..8

**Figure S5** Quantitative fluorescence microscopy for probe and ligand density quantification………..9

**Figure S6** Exclusion of DOTS from the cell spreading area..……………………………………..……10

**Figure S7** Quenching efficiency of DOTS……………………………………………………………..……11

**Figure S8** FLIM data showed DOTS eliminated intermolecular FRET at TCR clusters…………..……12

**Table S1** Table S1 List of DNA strands………………………………………………………………..….25

**Table S2** List of staple strands for DOTS in SLB experiments……………..………………….……….27

**Table S3** List of staple strands for DOTS in cell-cell experiments …….…………………………….....31

**Supplementary note 1** FLIM fitting and thresholding……………………………………...…………..11

**Supplementary note 2** oxDNA simulation parameters ……………….………………………....…17

**Supplementary note 3** Design of DNA hairpin tension sensors………………………………...…24

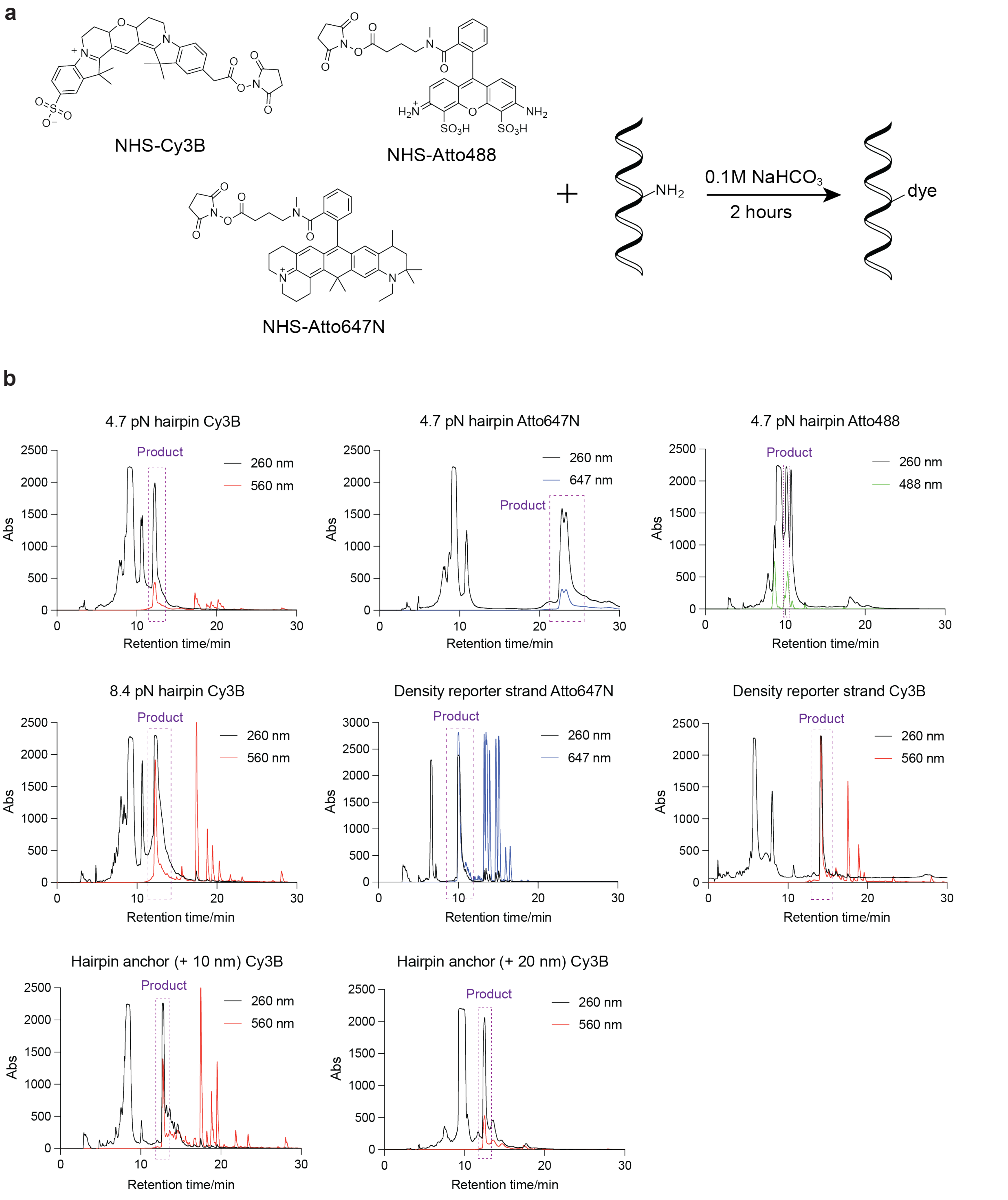

**Figure S1. Synthesis and purification of dye labeled oligonucleotides. a.** Reaction scheme for conjugation of amine-modified oligonucleotides with NHS-dyes **b.** HPLC traces of the dye-labeled targets. The dashed box represents the collected products which were validated by mass spectrometry.

#### 4.7 pN hairpin Cy3B

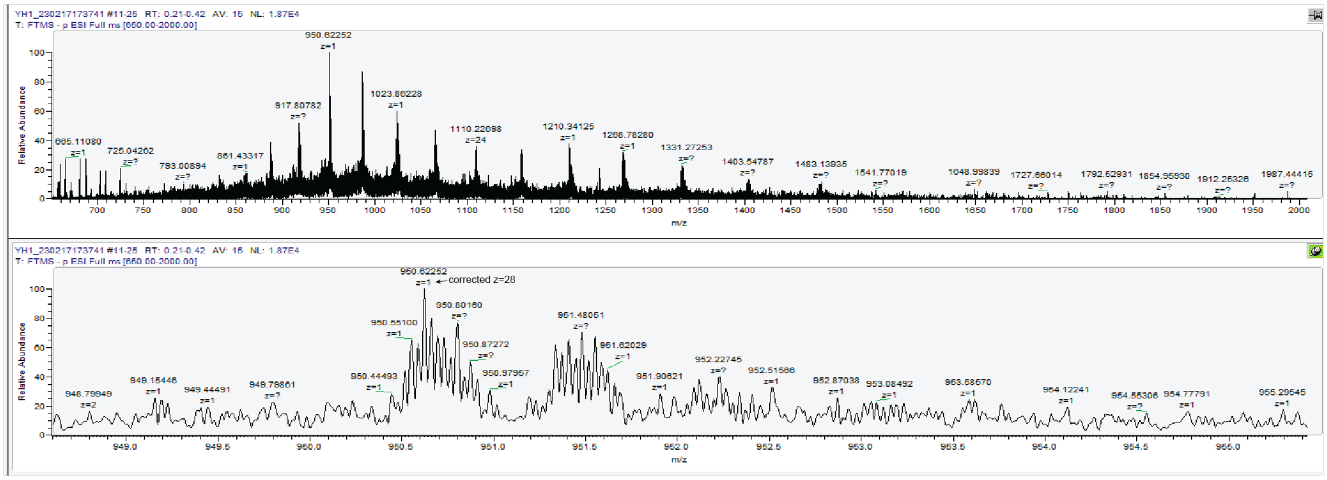

#### 4.7 pN hairpin Atto647N

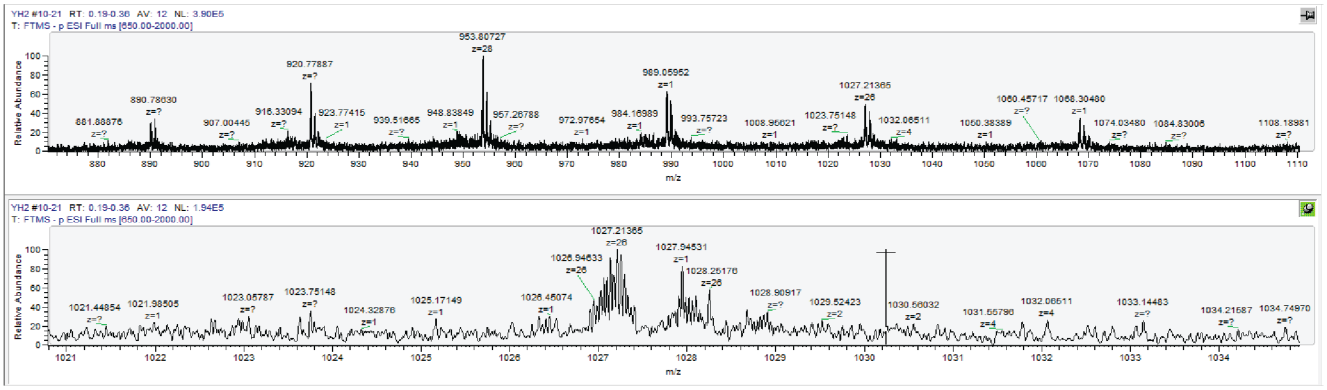

#### 4.7 pN hairpin Atto488

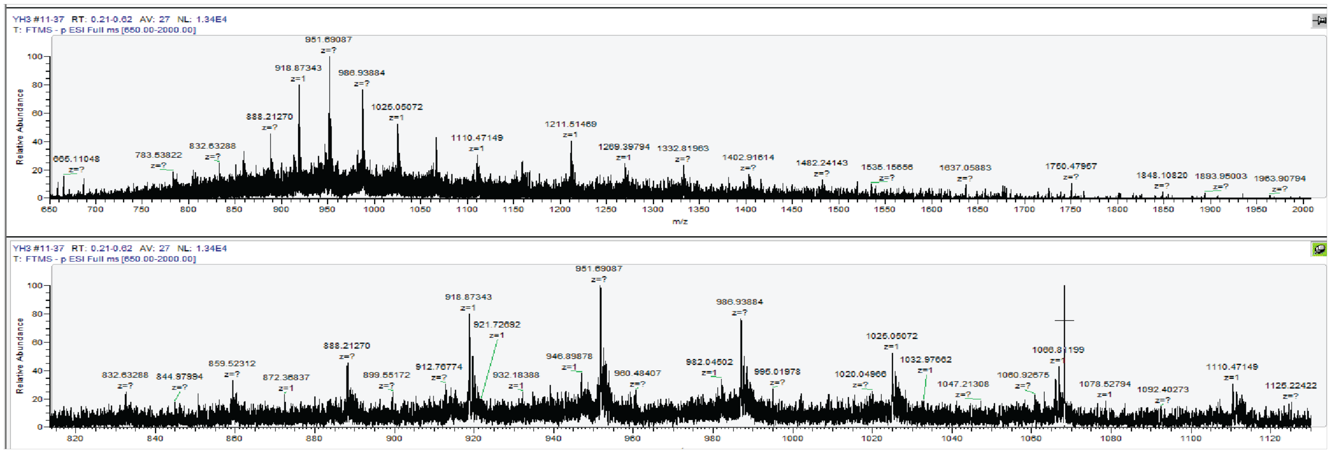

#### 8.4 pN hairpin Cy3B

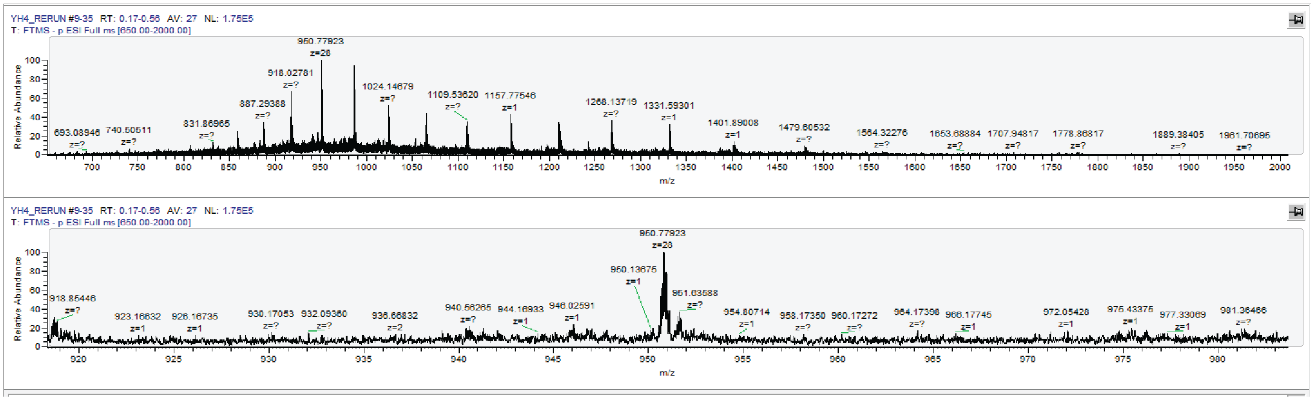

### Density reporter Atto647N

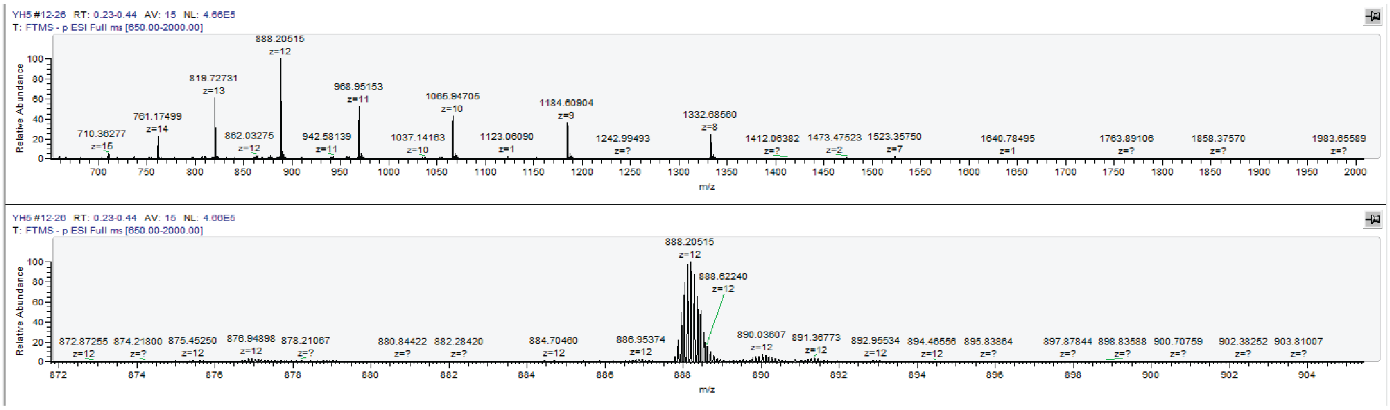

### Density reporter Cy3B

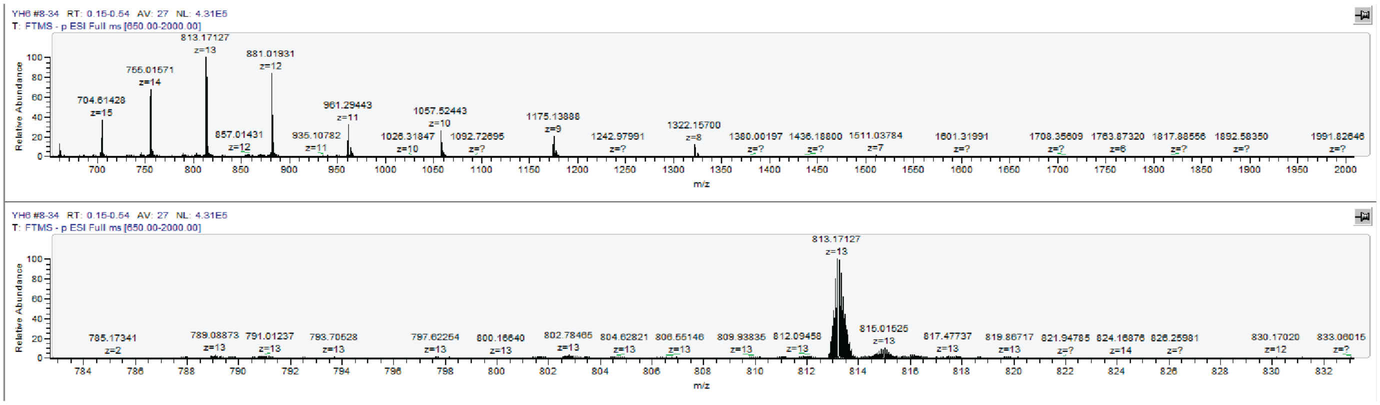

### Hairpin anchor (+ 10 nm) Cy3B

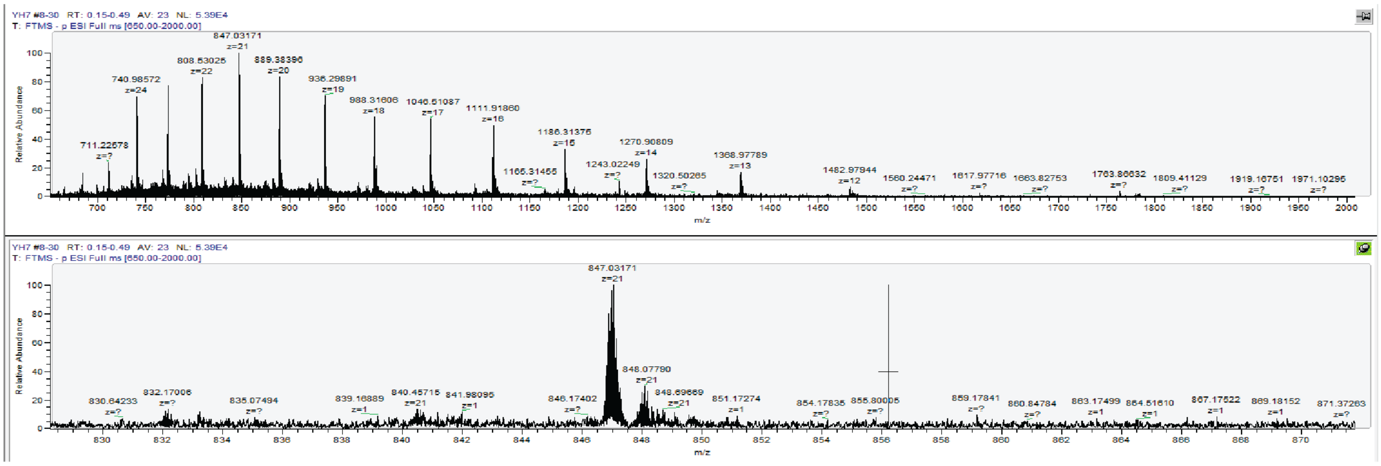

### Hairpin anchor (+ 20 nm) Cy3B

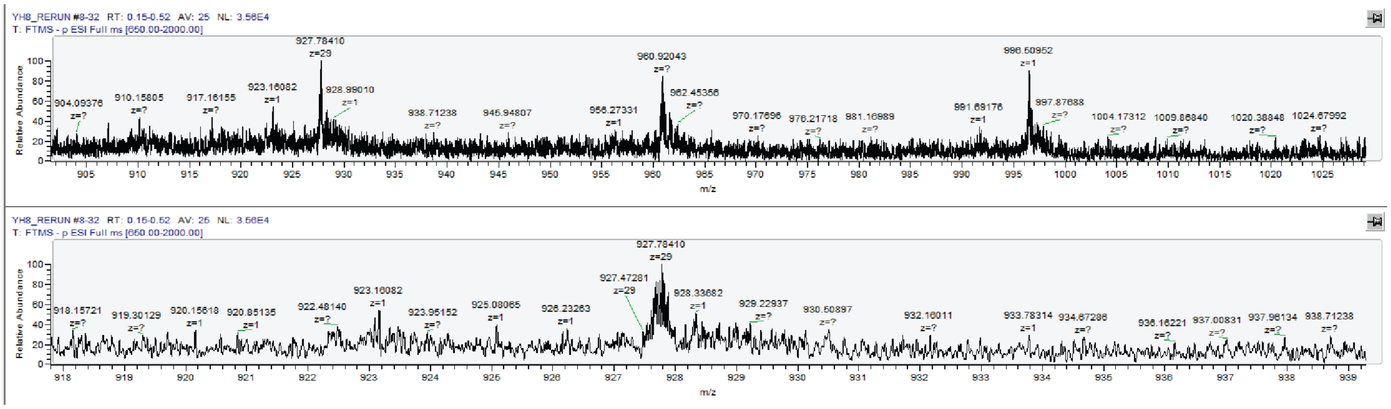

| Sample | Calculated mass | m/z found | Difference |
|---|---|---|---|
| 4.7 pN hairpin Cy3B | 26646.85 | 26646.43 | 0.42 |
| 4.7 pN hairpin Atto647N | 26733.08 | 26734.55 | 1.47. |
| 4.7 pN hairpin Atto488 | 26676.78 | 26676.34 | 0.44 |
| 8.4 pN hairpin Cy3B | 26651.65 | 26650.82 | 0.83 |
| Density reporter Atto647N | 10671.48 | 10671.46 | 0.02 |
| Density reporter Cy3B | 10585.25 | 10585.22 | 0.03 |
| Hairpin anchor (+ 10 nm) Cy3B | 17807.85 | 17809.67 | 1.82 |
| Hairpin anchor (+ 20 nm) Cy3B | 26933.75 | 26935.74 | 1.99 |

**Figure S2. Mass spectrometry characterization of dye labeled DNA oligos.** High resolution mass spectra of dye labeled oligos. Table shows calculated masses and measured masses.

**a**

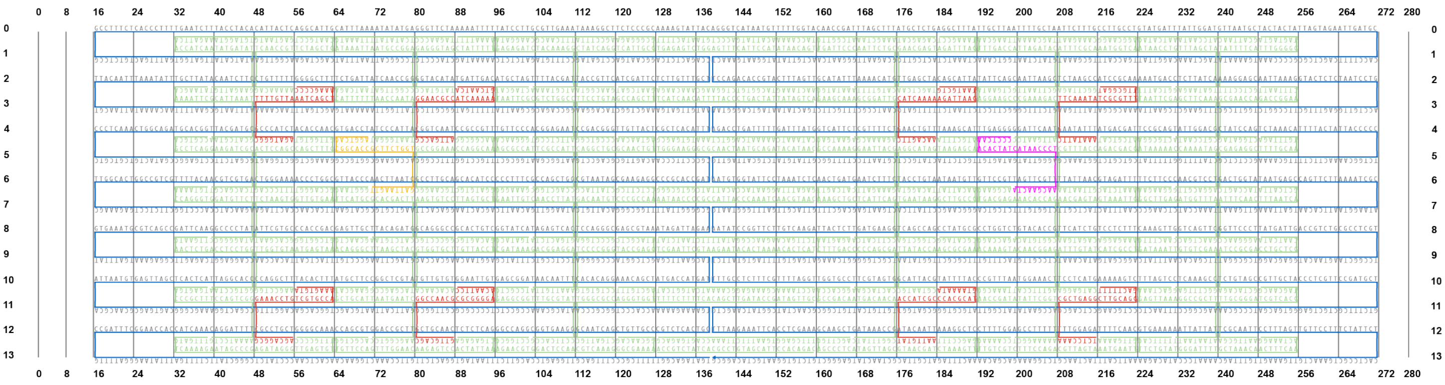

**b**

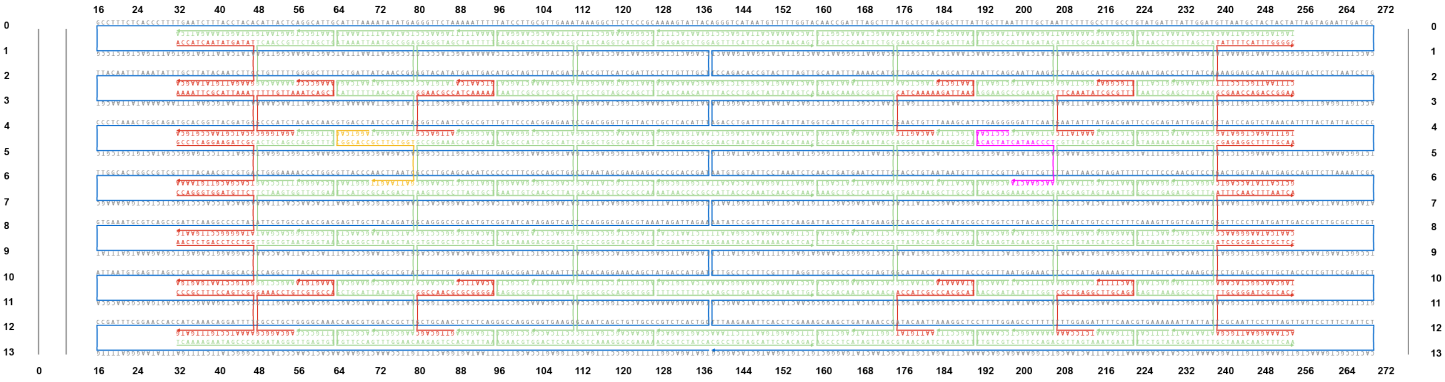

**Figure S3. Schematics of DNA origami structures used in this work. a** DNA origami was designed using CaDNAno based on the p7560 scaffold (blue). One staple strand (pink) was elongated at the 5’ end with the DNA hairpin sequence for visualizing TCR forces. One staple (orange) was labeled with Atto647N at the 3’ end to serve as a density reporter. Eight (8) staple strands (red) were elongated at the 3’ end with 21 base pairs that are complementary to DNA strand preinserted into SLBs to anchor DOTS to the SLB. **b** DNA origami design for cell-cell force experiment. 20 staple strands (red) were elongated at the 3’ end with 21 base pairs that are complementary to DNA strand on the cell membrane to attach DOTS to cell membrane.

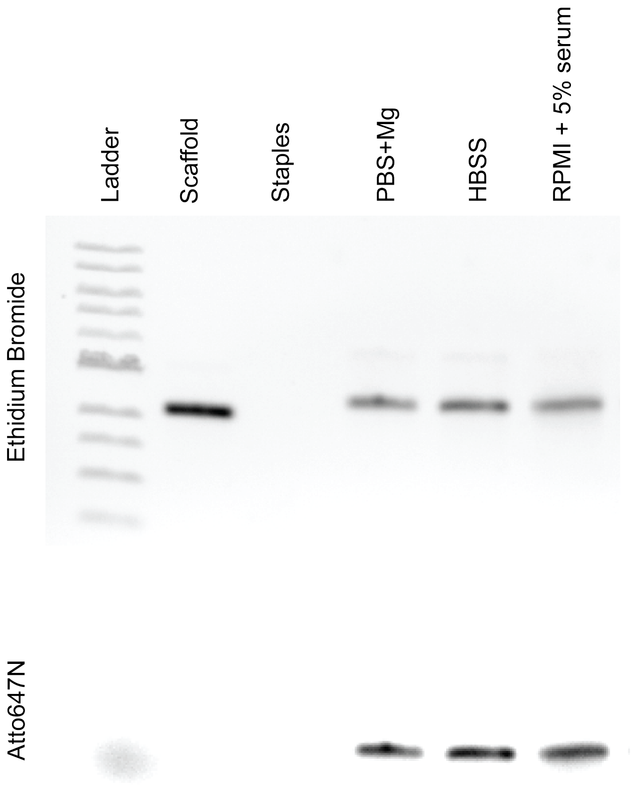

**Figure S4. Stability analysis of DNA origami under different cell imaging conditions.** DNA origamis were incubated in different imaging buffers/media for 1 hour at room temperature and subjected to gel electrophoresis. Overall structure stability and overhang stability were confirmed via Ethidium Bromide and Atto647N signal, respectively. No degradation was observed under all the imaging conditions.

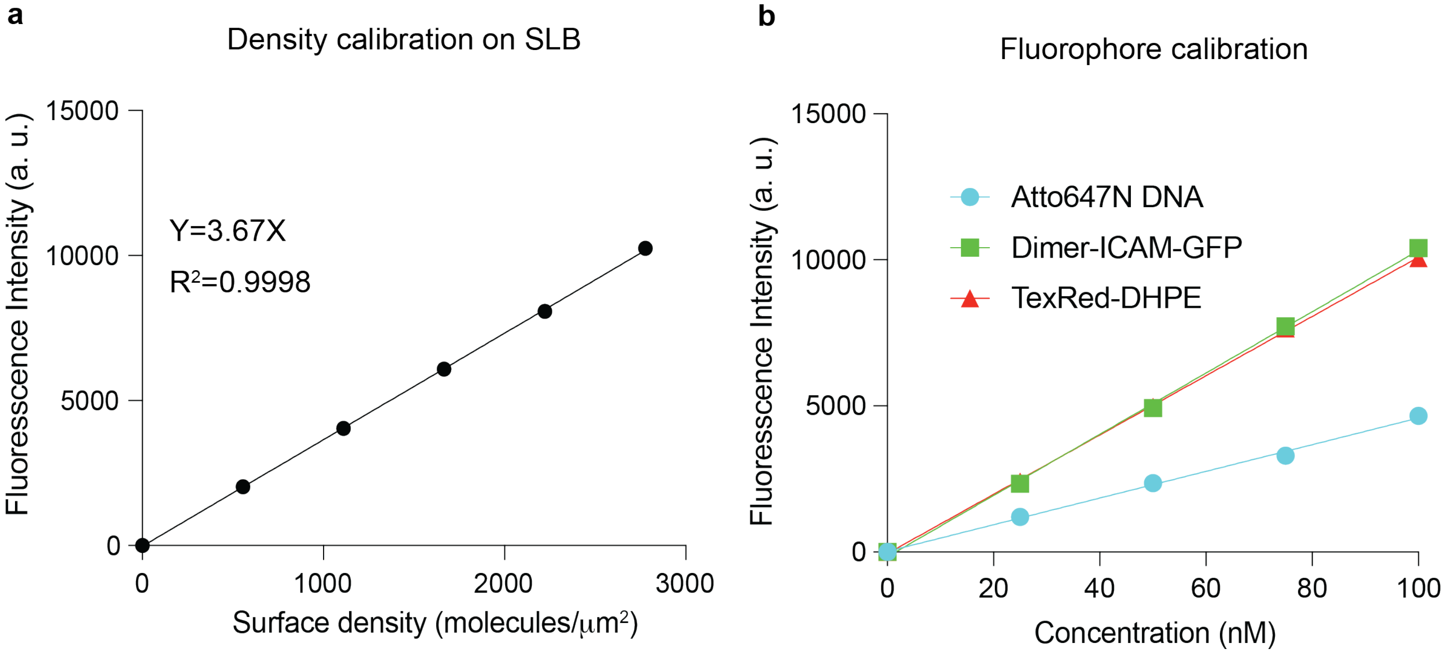

**c**

$$F\ (\text{factor}) = \frac{\text{Slope}_{\text{sample}}}{\text{Slope}_{\text{TR-DHPE}}} \qquad \text{Surface density} = \frac{\text{F. I.}_{\text{sample}}}{3.67 * F\ (\text{factor})}$$

**Figure S5 Quantitative fluorescence microscopy to determine densities of DOTS and ICAM-1 ligand on the SLB surface. a** TR-DHPE bilayer fluorescence calibration curve with known molecular densities. The intensity was obtained by averaging at least 6 images from each surface. Error bars indicate the SD which was smaller than the size of data symbol. **b** Plot showing the fluorescence intensities across various concentrations of fluorophores. Since sample fluorophore and Texas Red fluorophore on the lipid have different absorption and emission characteristics, the fluorescence intensity of the sample fluorophore needs to be calibrated to bilayer standard fluorophore. To obtain the F Factor, sample fluorescence and TR-DHPE lipid fluorescence at different concentrations were measured on a microscope. The F Factor was generated by dividing the slope of sample to that of TR-DHPE. **c** Equations used to calculate F factor and surface molecule densities.

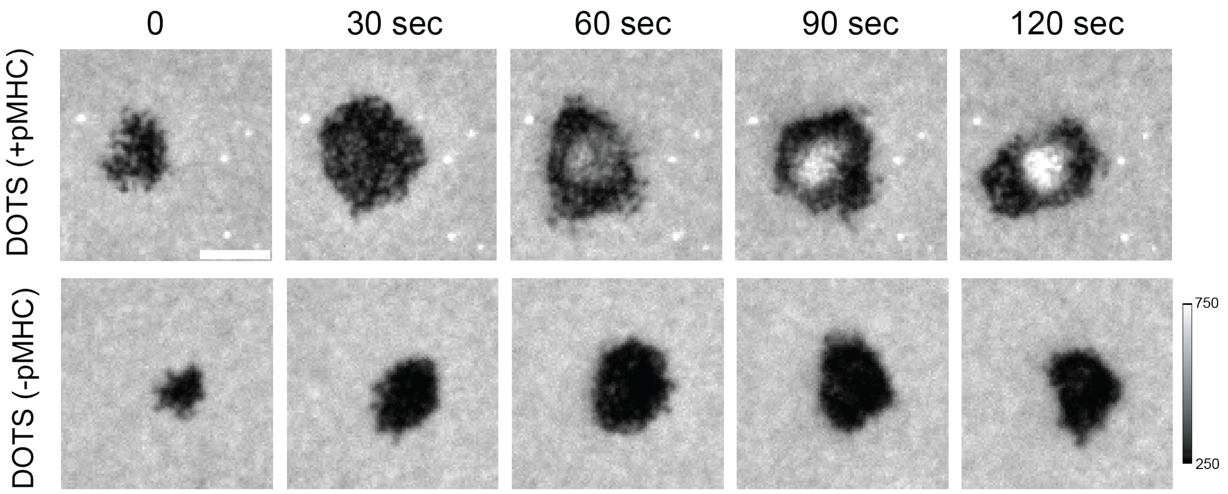

**Figure S6. Exclusion of DOTS from the cell spreading area.** Representative time lapse images showing the exclusion and clustering of DOTS in the T cell/SLB junctions. DOTS lacking antigen showed comparable levels of exclusion but did not exhibit any accumulation and clustering under T cells.

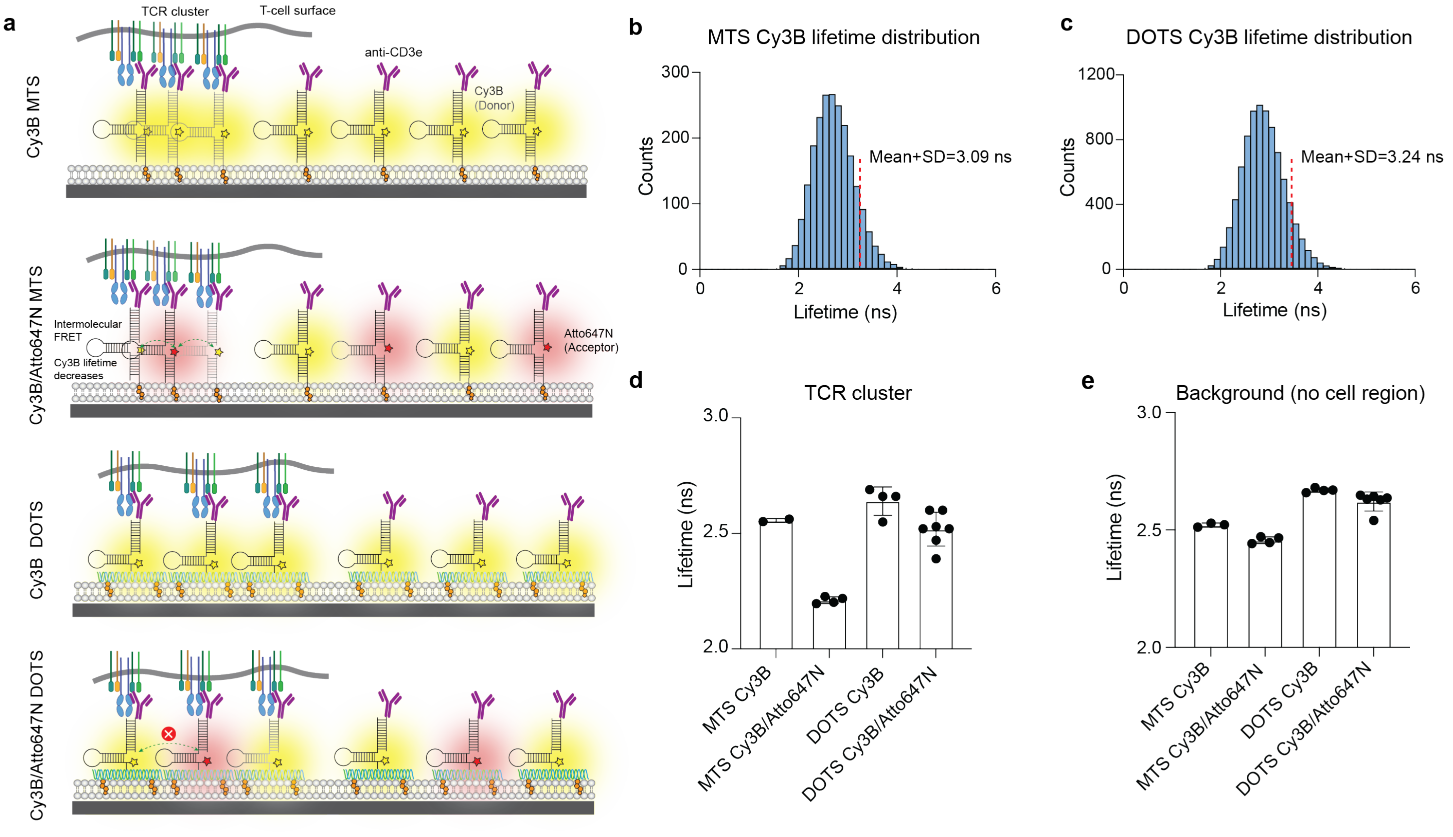

**Figure S7. FLIM data showed that DOTS eliminated intermolecular FRET between tension sensors at TCR clusters.** **a** Schematics showing SLBs coated with DOTS or MTS. **b** Fluorescence lifetime histogram of pixels on SLB surfaces containing only Cy3B MTS. **c** Fluorescence lifetime of pixels on SLB surfaces containing only Cy3B DOTS. Red dash lines indicate average lifetime + 1SD. **d** Bar graphs quantifying the average lifetimes of pixels in TCR clusters formed on Cy3B only MTS, Cy3B&Atto647N MTS, Cy3B only DOTS and Cy3B&Atto647N DOTS surfaces. **e** Bar graphs quantifying the average lifetimes of pixels at no cell region on Cy3B only MTS, Cy3B&Atto647N MTS, Cy3B only DOTS and Cy3B&Atto647N DOTS surfaces.

**Supplementary note 1: FLIM fitting and thresholding.** Fluorescence lifetime imaging microscopy (FLIM) requires data fitting to exponential decay curves, typically with a reconvolution decay model involving the instrument response function (IRF). A benefit of FLIM is that both spatial and temporal information are recorded; however, this creates a challenge as data is recorded separately within pixels rather than as an aggregate. From prior experience and reports from our lab,[1] we observe that pixels containing low photon counts (<25 photon/px) are noisy and can contain unreasonable lifetime values, both high and low. This can be avoided with spatial pixel binning or by thresholding pixels during analysis. To maintain spatial resolution and address this challenge through thresholding, we measured the lifetime values from surfaces containing only donor fluorescence (Cy3B) without cells. This is assumed to be the maximum theoretical lifetime. Data was fitted to a histogram and the average maximum lifetime + 1 SD was recorded (**Fig. S7b** and **Fig. S7c**). Note that SD refers to histogram width. This determined the lifetime cut-off as values above the lifetime max + 1SD are assumed to be noise (3.09 ns and 3.24 ns for hairpin and DOTS surfaces respectively). We then measured the mean photon count of pixels with lifetimes above the lifetime cut-off and determined that pixels under 25 photons should be excluded from analysis (SNR <5). FLIM fitting was conducted using both lifetime thresholding (<3.09 ns or 3.24 ns) and photon count thresholding (SNR > 5).

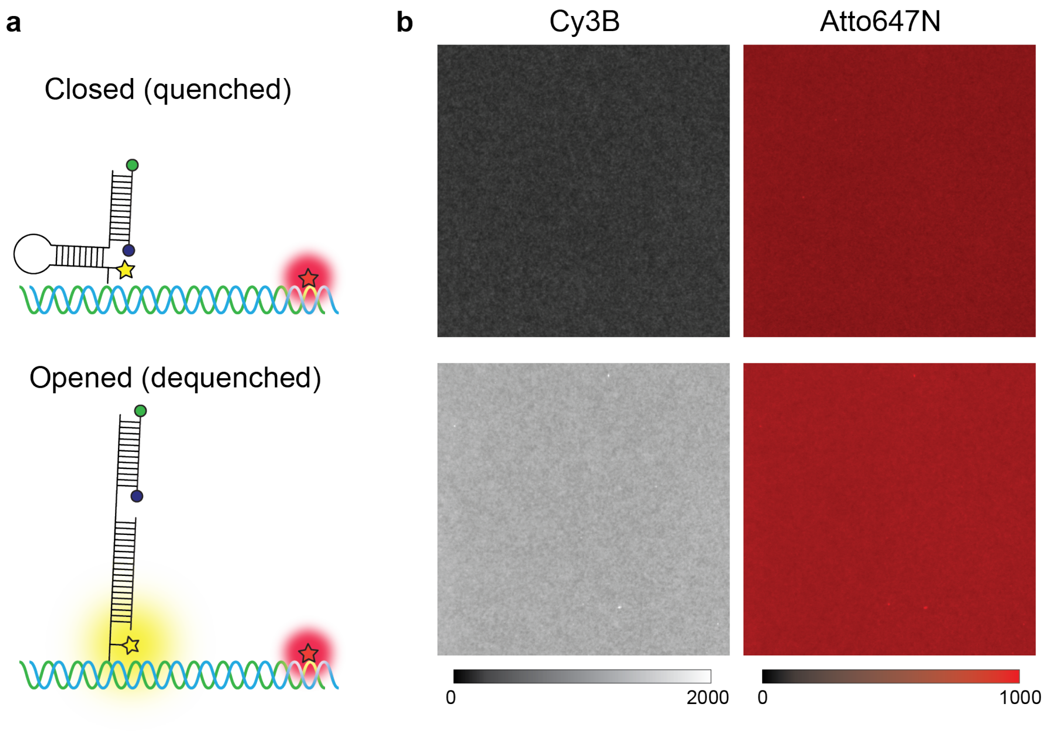

$$\text{Quenching effiency (\%)} = (1 - \frac{\text{Cy3B}_{\text{quenched}}}{\text{Cy3B}_{\text{dequenched}}} \times \frac{\text{Atto647N}_{\text{dequenched}}}{\text{Atto647N}_{\text{quenched}}}) \times 100\% = 69.4\%$$

**Figure S8. Quenching efficiency of DOTS. a** Schematic showing "closed" and "opened" hairpin probes on SLB surface. **b** Representative images showing the fluorescence of Cy3B and Atto647N of opened and closed DOTS. Quenching efficiency was calculated by dividing quenched Cy3B and dequenched Cy3B intensities normalized to Att647N intensities.

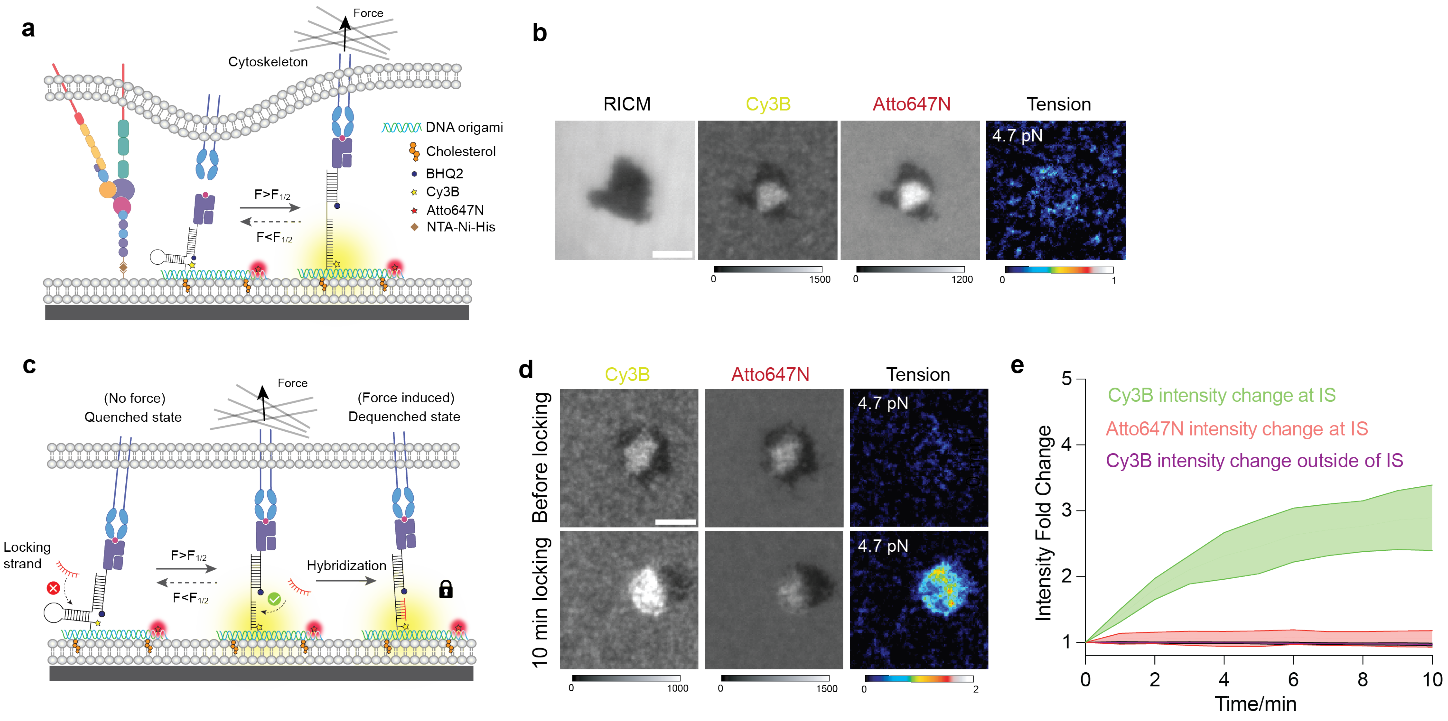

**Figure S9. Locking strategy specifically amplified tension signal at the immune synapse. a** Schematic showing that DOTS engages TCR and reports TCR force with a fluorescence increase in the Cy3B channel. **b** Representative microscope images showing T-cell spread on DOTS-SLB surface, but tension signal was not detectable. **c** Schematic showing that locking strand binds to mechanically unfolded hairpin and locks the hairpin in unfolded state to capture transient TCR force signal. **d** Representative images showing the addition of locking strand increased Cy3B fluorescence at the immune synapse, but Atto647N remained unchanged. **e** Plot showing the fluorescence increase after adding locking strand. At the immune synapse, Cy3B fluorescence increased up to 3-fold but Atto647N stayed constant. Cy3B fluorescence of ROI lacking cells did not change after addition of locking strand, confirming locking strand did not nonspecifically open hairpin. Data were obtained from >30 cells.

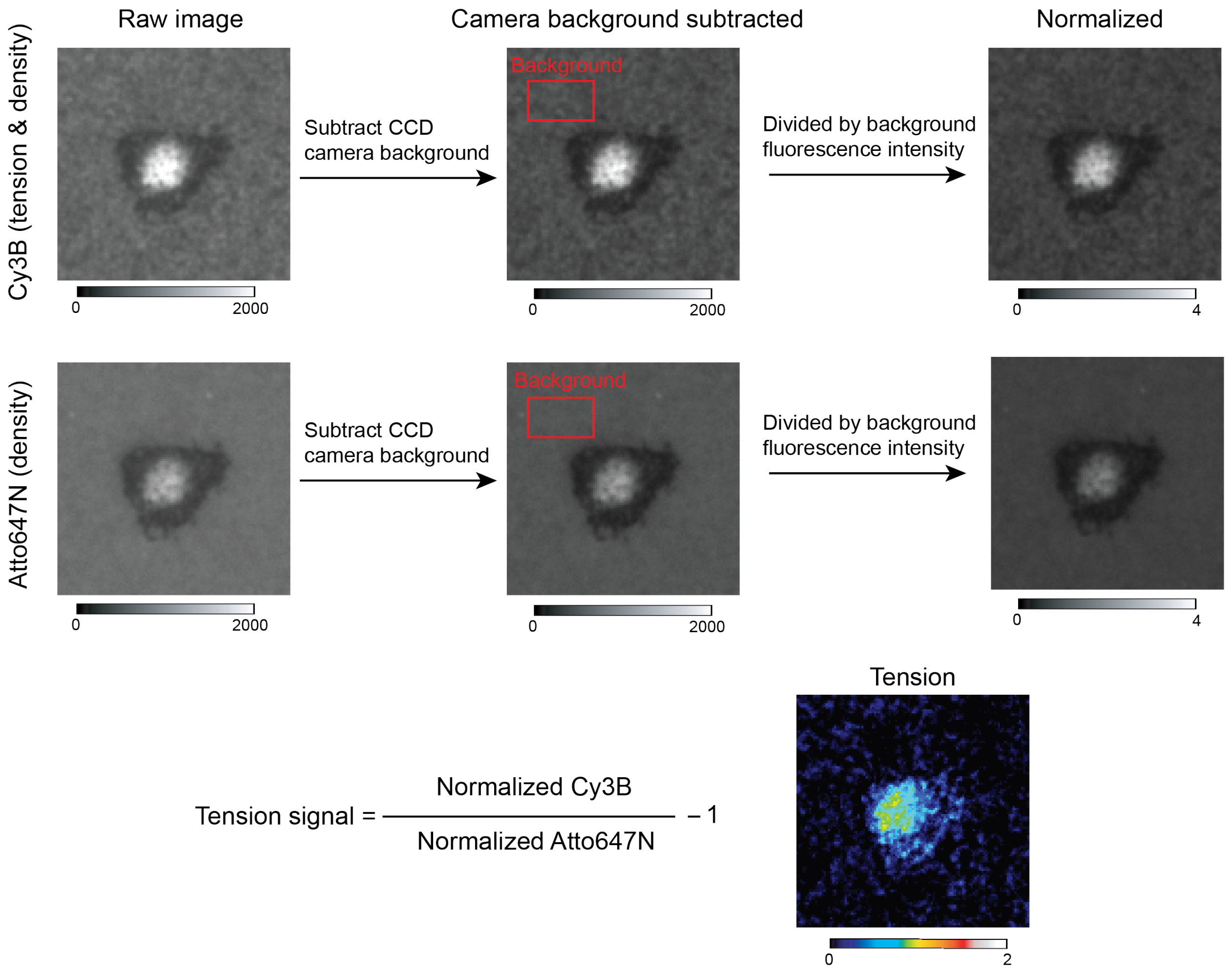

**Figure S10. Ratiometric analysis of TCR force signal.** To obtain tension signal, a series of image analysis was performed to extract tension signal from raw fluorescence images. First, raw Cy3B (signal includes both tension and density) and Atto647N (signal only includes density) fluorescence images were subtracted from EMCCD background of 200 a.u.. Afterwards, the images were normalized to background which was calculated by averaging ROIs lacking cells. Then the normalized Cy3B and Atto647N images were converted to tension signal by dividing Cy3B image by Atto647N image and subtracting the resulting image by 1.

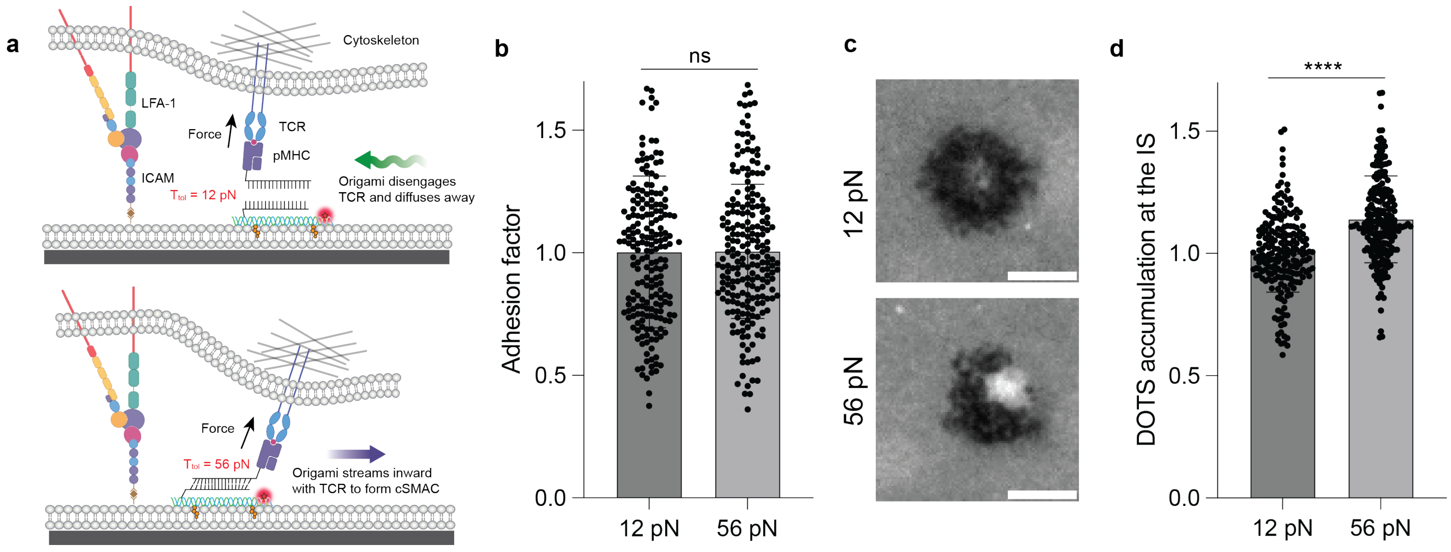

**Figure S11. TGT demonstrated that a subset of TCRs experience force > 12 pN. a** Schematic showing that TCR force ruptures TGT with a low force threshold (12 pN), and ruptured origami diffuses out of the immune synapse. By contrast, TGT with a high threshold (56 pN) is not ruptured by TCR force while translocating to the center of immune synapse. **b** Plot quantifying the area of immune synapse/adhesion zone formed on 12 and 56 pN TGT-DOTS surfaces. No statistical significance was observed. n>200 cells from three independent experiments and three independent mice. **c** Representative images showing the accumulation of DOTS signal at cSMAC on 12 and 56 pN TGT. **d** Plot quantifying the accumulation level of DNA origami at the immune synapse formed on 12 and 56 pN surfaces. **** p<0.0001

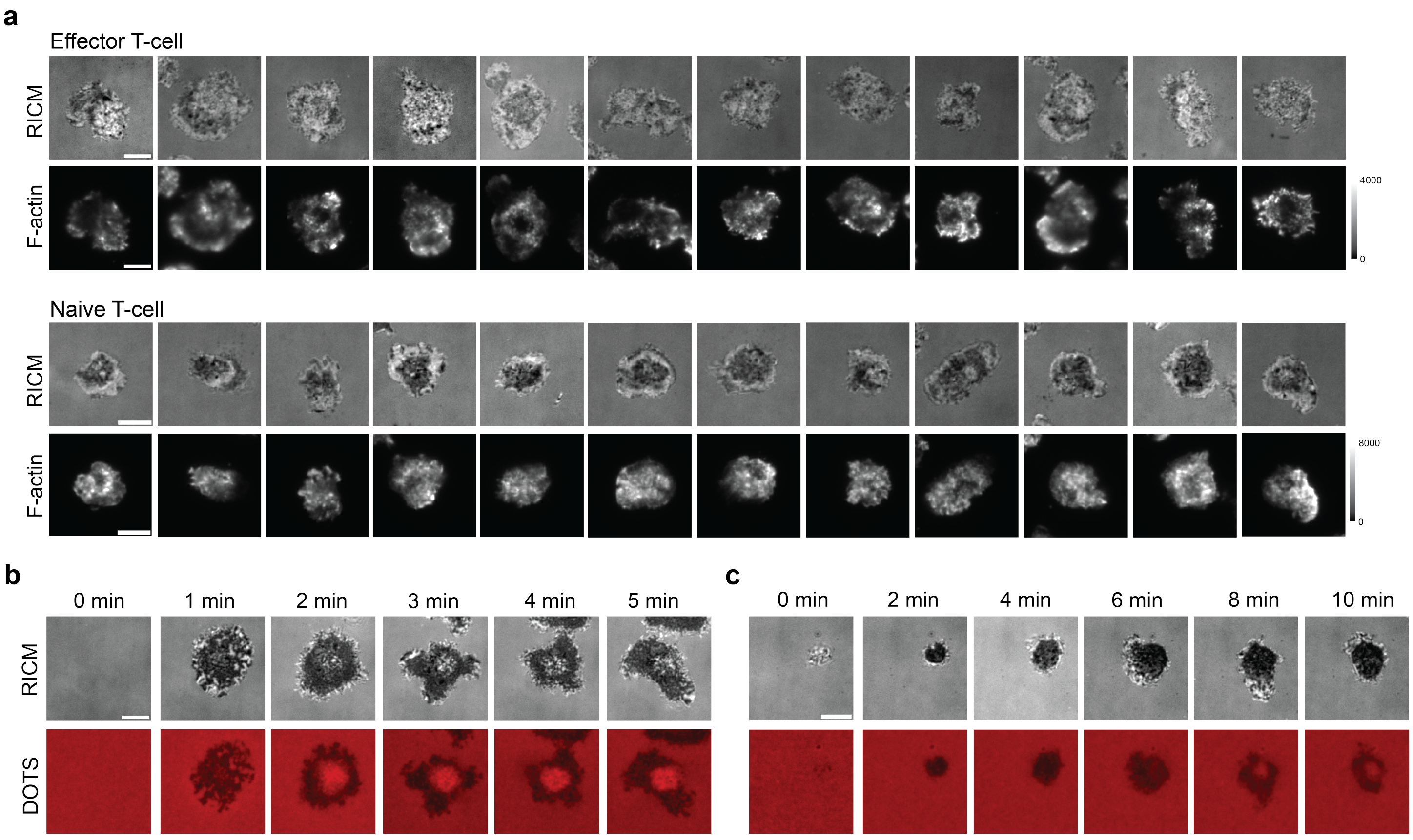

**Figure S12. F-actin dynamics of naïve and effector T-cell at the T-cell-SLB junctions a** Representative F-actin staining images showing the actin distribution of naïve T-cell and effector T-cell after 30 min spreading on SLB surfaces. Effector T-cell F-actin distribution was more heterogenous with a pronounced clearance at the cSMAC region. In contrast, naïve T-cell F-actin was more evenly and tightly distributed in the spreading area. **b-c** Representative time lapse images showing the formation of immune synapse of effector T-cell (b) and naïve T-cells (c) on SLB. Effector T-cell displayed a faster rate of immune synapse formation than that of naïve T-cell. Scale bars = 5 $\mu$m

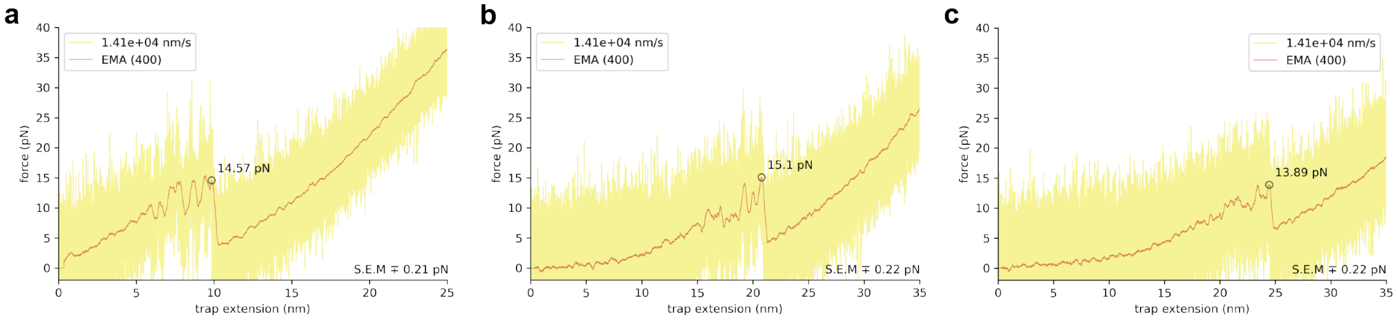

| DNA hairpin length | Original length | + 10 nm | + 20 nm |
|---|---|---|---|
| Replicate 1 (pN) | 14.57 | 10.35 | 13.89 |
| Replicate 2 (pN) | 12.52 | 15.1 | 14.82 |
| Replicate 3 (pN) | 12.27 | 12.06 | 13.69 |
| Average (pN) | 13.12 | 12.50 | 14.13 |

**Figure S13. Force threshold and height of DOTS are not correlated**. Force-extension curves of hairpin tension sensors on DOTS (**a** original length, **b** + 10 nm, **c** +20 nm) pulling at the rate of 1.4x104 nm/s in oxDNA simulations. The red curves indicate the smoothened data (400-point exponential moving average). The estimated force is marked at its peak in the graph along with its standard error of mean at the bottom right.

**Supplementary note 2:**

**DOTS height estimation:** The trajectories (oxDNA simulation timelapse) of the hairpin probes were imported into the oxView webserver. The transition point at which the structures start to unfold was noted and an earlier snapshot without the loss of any base pairing was used to measure heights. Since snapshots of the structure are printed in the simulation every millionth step ($10^6$), each frame in the simulation has a time interval of $10^6 \times 0.005 \times 3.03 \times 10^{-12}\, s = 15.15\, ns$. Hence, the height measurement was done ~15 ns before the start of hairpin rupture in the oxDNA simulation.

**oxDNA simulation parameters and conditions:** To validate that increasing linker length did not increase the force required to unfold DNA hairpin, we used oxDNA2 (version 2.4 published in June 2019) to simulate unfolding hairpins with different linker lengths (10.1063/1.4921957). We ran MD simulations on oxDNA on CPU to predict the rupture force of hairpins under force along the z-axis. The following parameters were used in the simulations:

backend = CPU
sim_type = MD
T = 37C
steps = 5e9
time_scale = linear
interaction_type = DNA2
use_average_seq = 1

verlet_skin = 0.05
salt_concentration = 0.156
thermostat = john
newtonian_steps = 103
diff_coeff = 2.5
dt = 0.005
print_conf_interval = 1e6

For all the simulations, harmonic traps of stiffness of 11.40 pN/nm were placed at the nucleotides of interest (i.e. the pMHC linked nucleotide and the nucleotide anchored to the origami). The effective trap stiffness can be calculated using the following equation: ${}^{1}/_{k_{eff}} = {}^{1}/_{k_1} + {}^{1}/_{k_2}$

For all the simulations, harmonic traps (springs in simulation) of stiffness of 11.40 pN/nm were placed at the nucleotides of interest (i.e. the pMHC linked nucleotide and the nucleotide anchored to the origami). The effective trap stiffness can be calculated using the following equation: ${}^{1}/_{k_{eff}} = {}^{1}/_{k_1} + {}^{1}/_{k_2}$. The ligand nucleotide traps were moved at a rate of $5 \times 10^{-8}$ (length per unit of time in oxDNA units) along the z-direction and when converted into SI units it yields a loading rate $= \frac{5 \times 10^{-8} \times 0.8518\ \text{nm}}{3.03 \times 10^{-12}\ s} = 1.4 \times 10^{4}\ nm/s$. To obtain the force-extension curves, we extracted the extensions of individual harmonic traps from the corresponding attached nucleotides and then projected it along the z-axis. The force is calculated by multiplying the total projected trap extension with $k_{eff}$. Combined trap extensions were used along with $k_{eff}$ to average random thermal fluctuations as previously done in literature (10.1021/acsnano.8b01844). The graph with $1.9\times10^5$ data points is then smoothed with a 400-point exponential moving average (EMA) of the data points using python (10.1038/s41586-020-2649-2). The peak rupture force was then estimated using SciPy find peaks module on the smoothened graph (10.1038/s41592-019-0686-2).

From the simulations, the rupture of hairpins remained relatively within the range of 10-14 pN and did not correlate with linker length. It must also be noted oxDNA runs have an inherent level of stochasticity (as seen in the replicates) that results in small variations in force estimation. Note that in oxDNA simulations we measure peak force as the maximum force before the hairpin unfolding transition, therefore this can lead to higher estimation of forces compared to the experimental calibration previously done using biomembrane force probe.[2] In a typical "force ramp" setup, the rupture force is highly dependent on the loading rate (i.e. the greater the loading rate the greater the rupture force).

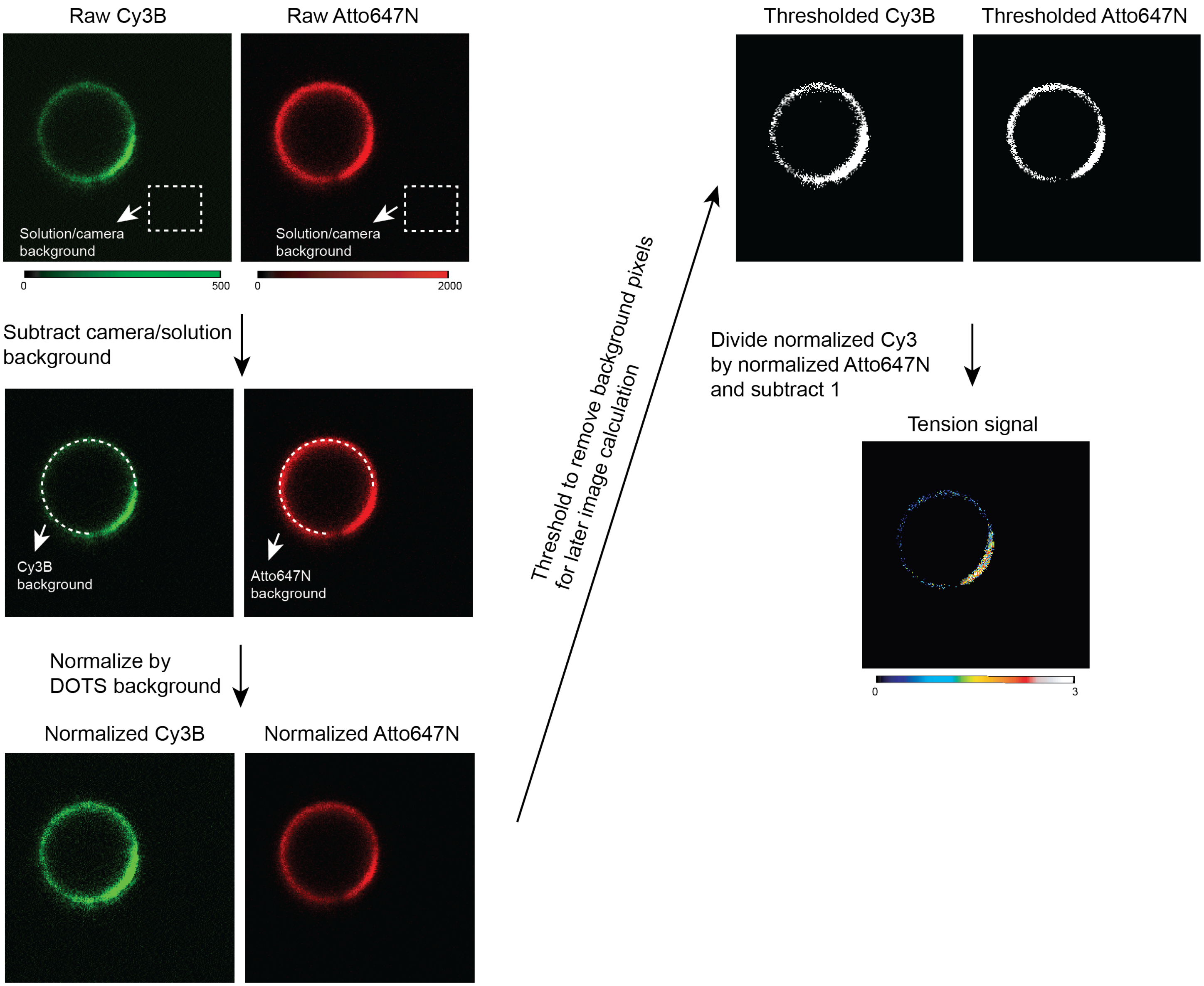

**Figure S14. TCR force analysis protocol for cell-SSLB experiment.** First, raw Cy3B (signal includes both tension and density) and Atto647N (signal only includes density) fluorescence images were subtracted from camera/solution background (intensity of regions lacking SSLB). Second, Cy3B and Atto647N images were normalized by a defined region not engaging cells. Third, because background subtracted images have zero values which would introduce infinite values/errors in following image calculations, a threshold mask was applied to exclusively select the pixels associated with the SSLB. Fourth, tension signal was obtained by dividing normalized Cy3B image (signal includes both tension and density) by Atto647N image (signal only includes density) and subtracting 1.

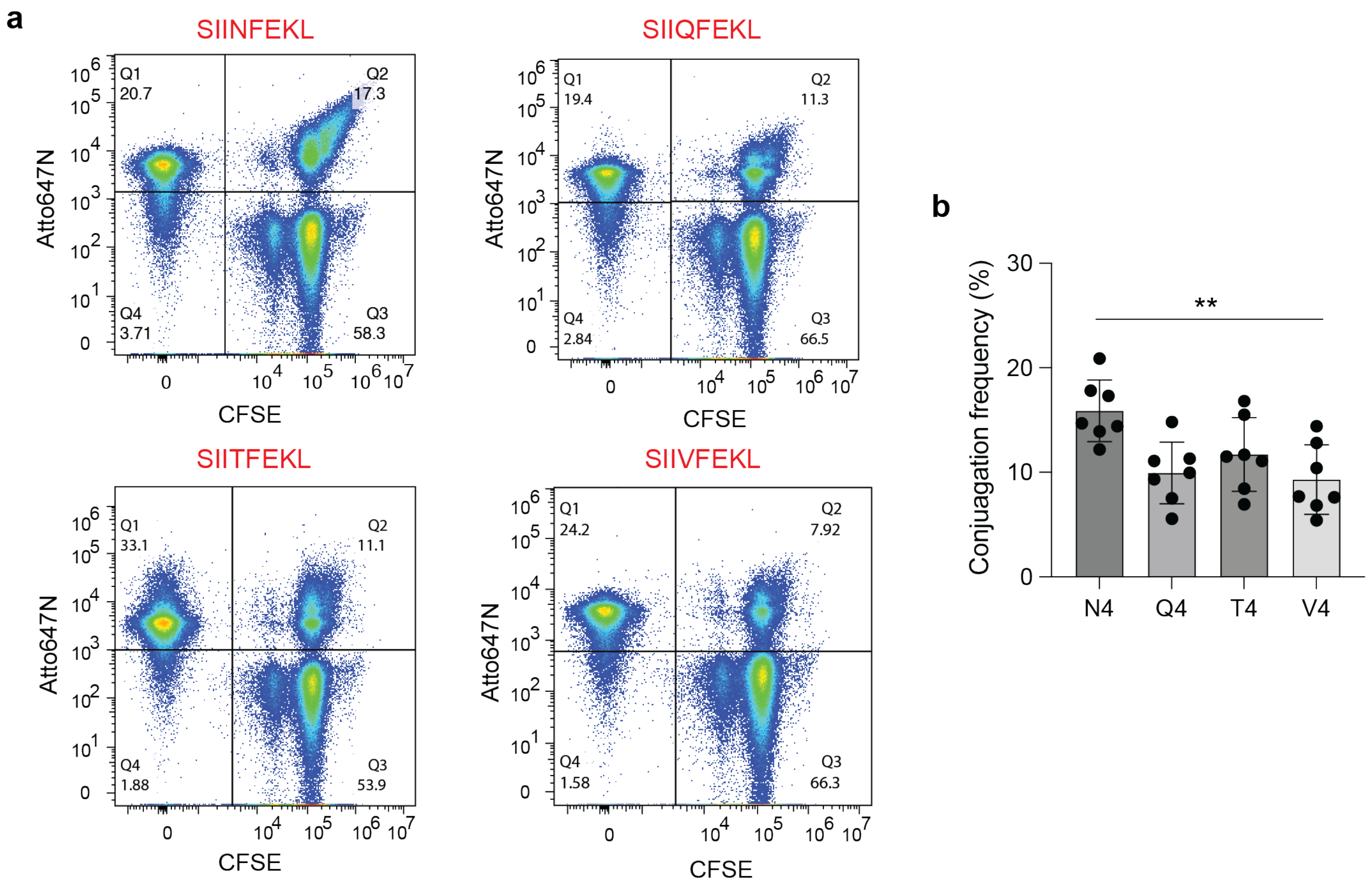

**Figure S15. Flow analysis of T-cell-SSLB conjugates under different antigen conditions. a** Flow fluorescence quadrant plots showing the T-cell (CFSE positive), SSLB (Atto647N positive) and T-cell-SSLB conjugate (Dual positive) populations. SSLB were coated with DOTS modified with different antigens. **b** Plot quantifying the frequency of conjugates. Conjugates formed more efficiently under agonist (N4) condition compared to antagonists. ** p= 0.0033, n= 6 replicates from 6 independent experiments.

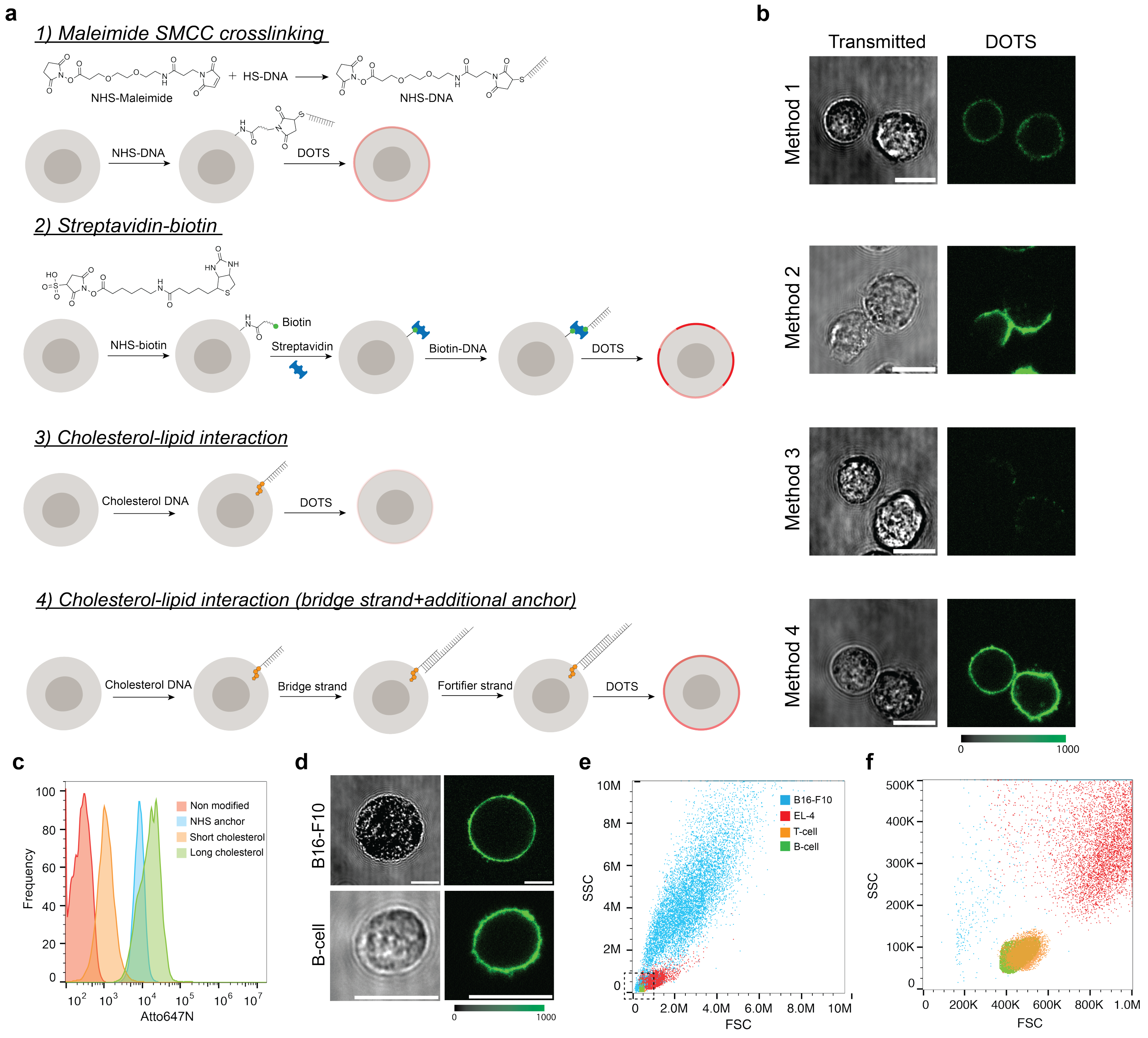

**Figure S16. Different strategies to functionalize cell membrane with DOTS. a** Schematics showing different chemistries used to modify cell membrane with DOTS. DNA strands complementary to the elongations on the DOTS were anchored onto cell membrane through thiol-maleimide reaction, streptavidin-biotin interaction or cholesterol lipid interaction. **b** Representative images showing the DOTS labeling intensities on EL-4 membrane with different functionalizing strategies. **c** Flow cytometry fluorescence histogram quantifying the labeling intensities under different functionalizing strategies. **d** Representative images showing DOTS signal on B16-F10 and resting B-cell membranes. DOTS were anchored through strategy 4 in panel a **e** Flow cytometer scatter plot showing the size distribution of different cell lines **f** Zoom-in scatter plot of selected ROI in **e** showing the resting T-cell and B-cell have comparable size. Scale bars = 10 μm

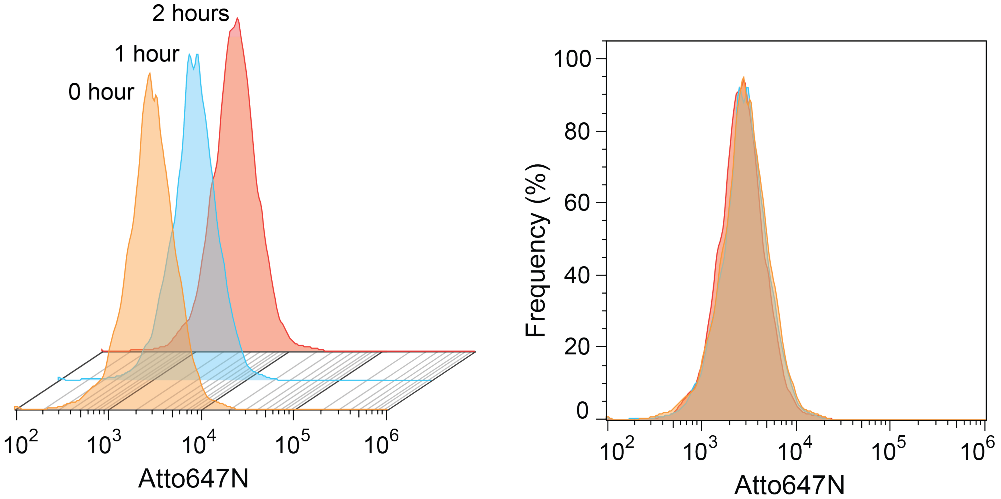

**Figure S17. Stability of DOTS on the B-cell membrane.** Flow fluorescence histograms of DOTS modified B-cells. No fluorescence change was observed within two hours at room temperature, indicating a strong stability of DOTS on B-cell membrane and low dissociation rate.

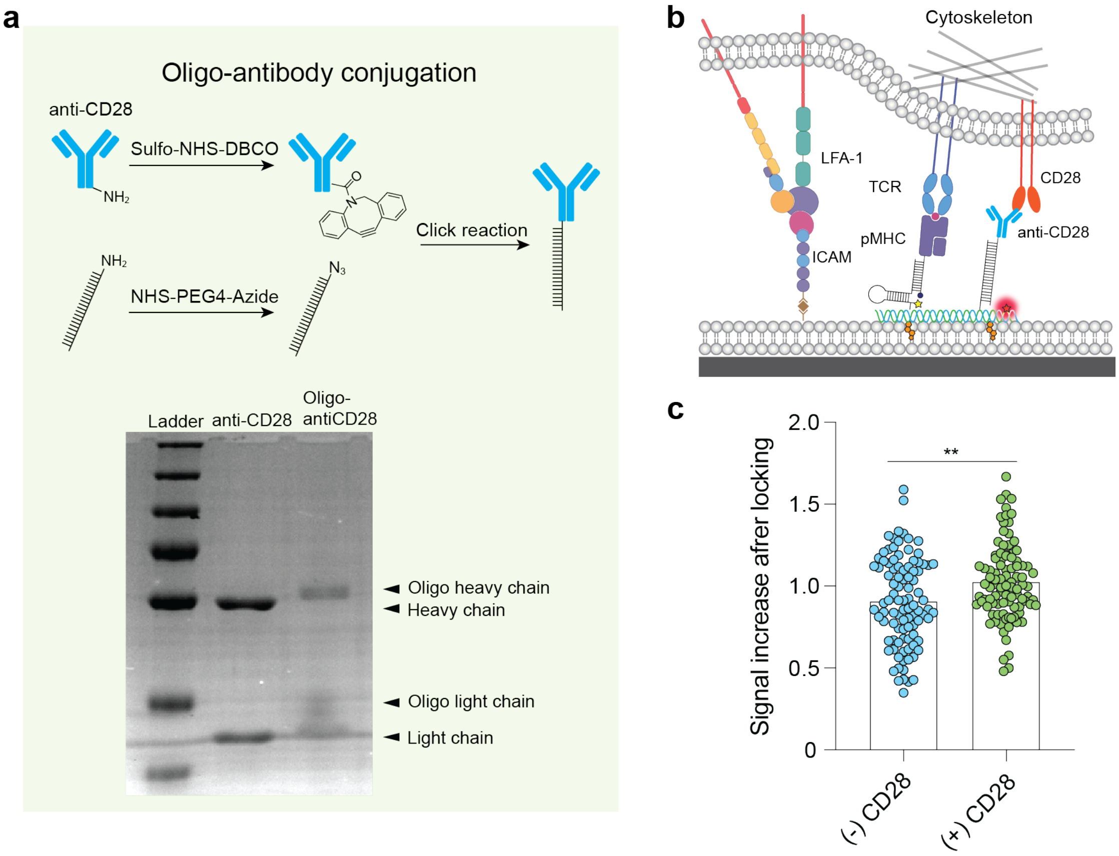

**Figure S18. CD28 engagement enhances TCR force signal. a** Schematic showing the workflow of conjugating anti-CD28 to DNA through copper free click reaction. Amine-DNA and Anti-CD28 was labeled using Azide-NHS and DBCO-NHS, respectively. Successful conjugation was confirmed using reducing SDS-PAGE. Arrows indicate anti-CD28 and DNA-anti-CD28 fragments. **b** Schematic showing the anti-CD28 was introduced to DNA origami platform through DNA hybridization. pMHC and anti-CD28 were on the same DOTS surrounded by ICAM-1 ligands. **c** Dot plot comparing the TCR force signals in the presence and absence of anti-CD28 on the DOTS. At least 95 cells from two independent mice were analyzed. ** p=0.0013

**Supplementary note 3: Design of DNA hairpin tension sensors**

For DNA hairpin structures, the free energy change during transition from the folded state to the unfolded state under an applied F can be described as:

$$\Delta G = \Delta G_{\text{unfold}} + \Delta G_{\text{stretch}} - F\Delta x + k_B T ln\left(\frac{[unfolded]}{[folded]}\right) \qquad (1)$$

When F = $F_{1/2}$, at equilibrium (ΔG=0) the DNA hairpin molecule has an equal probability of being in folded and unfolded states such that ([unfolded])/([folded]) = 1. Accordingly, above equation can be reorganized as the following:

$$F_{1/2} = \frac{(\Delta G_{\text{unfold}} + \Delta G_{\text{stretch}})}{\Delta x} \qquad (2)$$

where, $\Delta G_{unfold}$ is the free energy for unfolding the hairpin without force and opposite to $\Delta G_{fold.}$ $\Delta G_{fold}$ was acquired from IDT oligoanalyzer 3.1, which uses the UNAfold software package to calculate $\Delta G_{fold}$ under T-cell imaging condition (25 °C, 140 mM monovalent salt and 2 mM divalent salt). $\Delta G_{stretch}$ is the free energy for stretching the ssDNA from its folded coordinates, which can be calculated with the worm like chain model below:

$$\Delta G_{\text{stretch}} = \left(\frac{k_B T}{L_p}\right)\left[\frac{L_0}{4\left(\frac{1-x}{L_0}\right)}\right]\left[3\left(\frac{x}{L_0}\right)^2 - 2\left(\frac{x}{L_0}\right)^3\right] \qquad (3)$$

where Lp is the persistence length of ssDNA (~1.3 nm), $L_0$ is the contour length of ssDNA (0.63 nm per nucleotide), x is the hairpin extension from equilibrium and can be calculated by using (0.44*(n-1)) nm, where n represents the number of the bases comprising the stem-loop of hairpin. Note we subtract a distance of 2 nm from x to get Δx in equation 2 because initial separation between the stem termini is set by the diameter of the hairpin stem duplex (effective helix width = 2 nm).

The parameters and $F_{1/2}$ of hairpins with 22% and 77 GC% content were listed below:

| Name | 4.7 pN hairpin | 8.4 pN hairpin | 13.1 pN hairpin |
|---|---|---|---|
| Stem loop sequence | GTA TAA ATG TTT TTT TCA TTT ATA C | GGG GAG GAG TTT TTT TCT CCT CCC C | GCG CGC GCG CGC TTT TGC GCG CGC GCG C |
| Nucleotide number | 25 | 25 | 28 |
| GC content | 22% | 77% | 100% |
| Δx (nm) | 8.56 | 8.56 | 9.88 |
| $\Delta G_{unfold}$ (kJ/mole) | 24.23 | 43.60 | 102.17 |
| $\Delta G_{stretch}$ (kJ/mole) | 16.99 | 16.99 | 19.28 |
| Calculated $F_{1/2}$ (pN) | 7.99 | 11.7 | 20.42 |
| Calibrated $F_{1/2}$ (pN) | 4.7 | 8.4 | 13.1 |
| Correction Factor f | 0.41 | 0.39 | 0.36 |

The $F_{1/2}$ s of "4.7 pN hairpin" and "13.1 pN" were previously calibrated with biomembrane force probe under the T-cell imaging condition: 25 °C, 140.6 mM $Na^+$ and 1.8 mM $Mg^{2+}$.[2] By comparing the calibrated $F_{1/2}$ values of the 4.7 pN and 13.1 pN probes to their calculated $F_{1/2}$ values, we determined a correction factor f by using the following equation:

$$f = \frac{\left(F_{1/2\ \text{Calculated}} - F_{1/2\ \text{Calibrated}}\right)}{F_{1/2\ \text{Calibrated}}}$$

By averaging the factor of 4.7 and 13.1 pN hairpins, we obtained a generalized f factor which is further used to determine the calibrated $F_{1/2}$ of "8.4 pN hairpin".

**Table S1 List of DNA strands**
All oligonucleotides were custom synthesized by Integrated DNA Technologies (Coralville, IA), except for the BHQ1 strand, which was synthesized by Biosearch Technologies (Novato, CA). Table S2 includes the names and sequences for all oligonucleotides used in this work except for DNA origami staples. The colors of the highlighted sequences match the colors in Figure S3. Structures of the modifications are shown in Figure S1.

| Name | Description | Sequence (5' to 3') |
|---|---|---|
| 4.7 pN hairpin | DNA origami staple with stem-loop hairpin ($F_{1/2}$=4.7 pN) and overhang complementary to ligand strand | CCC TCA AAC ACT ATC ATA ACC CTA ACG AAC TA/iAmMC6T/ TTT GTA TAA ATG TTT TTT TCA TTT ATA CTT TGT GTC GTG CCT CCG TGC TGT G |
| Ligand strand | Strand carrying antigen and quencher | /5_Biotin/ CAC AGC ACG GAG GCA CGA CAC /3_BHQ2/ |
| 4.7 pN locking strand | Strand is partially complementary to 4.7 pN hairpin | AAA AAA CAT TTA TAC |
| Fully complementary strand | Strand is fully complementary to 4.7 pN hairpin | GTA TAA ATG AAA AAA ACA TTT ATA C |
| Density reporter strand | DNA origami staple labeled with fluorophore as a density reporter | AGG TCA CCG GCA CCG CTT CTG GTG ATT AAG TT/3AmMO/ |
| 8.4 pN hairpin | DNA origami staple with stem-loop hairpin ($F_{1/2}$=8.4 pN) and overhang complementary to ligand strand | CCC TCA AAC ACT ATC ATA ACC CTA ACG AAC TA/iAmMC6T/ TTT GGG GAG GAG TTT TTT TCT CCT CCC CTT TGT GTC GTG CCT CCG TGC TGT G |
| 8.4 pN locking strand | Strand is partially complementary to 4.7 pN hairpin | AAA AAA CTC CTC CCC |
| TGT bottom strand | DNA origami staple with strand complementary to TGT top ligand strand | CCC TCA AAC ACT ATC ATA ACC CTA ACG AAC TA TT T GTG AAA TAC CGC ACA GAT GCG |
| 12 pN TGT top strand | 12 pN TGT top strand carrying antigen next to DNA origami anchor site (unzipping geometry) | CGC ATC TGT GCG GTA TTT CAC /Biotin/ |
| 56 pN TGT top strand | TGT top strand carrying antigen at opposite terminal to DNA origami anchor site (shearing geometry) | CGC ATC TGT GCG GTA TTT CAC /Biotin/ |
| Hairpin anchor strand (+ 10 nm) | DNA origami staple with elongation complementary to the conventional hairpin strand | CCC TCA AAC ACT ATC ATA ACC CTA ACG AAC TA T TTG CTG GGC TAC GTG GCG CTC TT /3AmMO/ |
| Hairpin anchor strand (+ 20 nm) | DNA origami staple with elongation complementary to the conventional hairpin strand | CCC TCA AAC ACT ATC ATA ACC CTA ACG AAC TAT TTT TTT TTT TTT TTT TTT TTT TTT TTT TTT TTG CTG GGC TAC GTG GCG CTC TT/3AmMO/ |
| MTS 4.7 pN hairpin strand | Hairpin strand with stem-loop region ($F_{1/2}$=4.7 pN) and two arms that binds to the hairpin anchor strand and ligand strand for higher antigen | GTG AAA TAC CGC ACA GAT GCG TTT GTA TAA ATG TTT TTT TCA TTT ATA CTT TAA GAG CGC CAC GTA GCC CAG C |

| | | |
|---|---|---|
| Ligand strand for higher antigen | Strand carrying antigen and quencher | /5_BHQ1/ CGC ATC TGT GCG GTA TTT CAC /3_Biotin/ |
| Cholesterol SLB strand | Cholesterol SLB strand is complementary to elongations on the origami for anchoring origami onto SLB | GTT CGT CCG CTC GCC TGC TTG /3CholTEG/ |
| Second ligand anchor strand | DNA origami staple with elongation complementary to second ligand strand | AGG TCA CCG GCA CCG CTT CTG GTG ATT AAG TT TTT CAC TCC CGT CCA CAT TGC TAC TAC TAT CAT |
| Second ligand strand | DNA strand used to label anti-CD28 | /5AmMC6/ ATG ATA GTA GTA GCA ATG TGG ACG GGA GTG |
| Cholesterol cell membrane strand | Cholesterol SLB strand is used to tag cell membrane with DNA and is complementary to part of the bridge strand | GAT GAA TGG TGG GTG AGA GGC /CholTEG/ |
| Bridge strand | Strand that connects cholesterol cell membrane strand and elongations on the DOTS to anchor DOTS onto cell membrane | GCC TCT CAC CCA CCA TTC ATC TTT TTT TTT TTT TTT TTT GTT CGT CCG CTC GCC TGC TTG |
| Fortifier strand | Strand binds to the remaining portion of bridge strand to rigidify the bridge strand | AAA AAA AAA AAA AAA AAA |
| MTS ligand strand | Ligand strand of DNA hairpin directly anchored to SLB | /5_bitoin/TT TGC TGG GCT ACG TGG CGC TCT T/3AmMO/ |
| MTS anchor strand | Anchor strand of DNA hairpin directly anchored to SLB | CGC ATC TGT GCG GTA TTT CAC /3_CholTEG/ |

**Table S2. List of staple strands for DOTS in SLB experiments**

| Start 5' | End 3' | Note | Sequence |
|---|---|---|---|
| 6[198] | 8[199] | Staple | ACGGAACTGACGAGAAACACCAGCGGTGTACA |
| 4[38] | 6[39] | Staple | CCGTGCAGCCTCAGGAAGATCGCTCACGACGT |
| 8[215] | 6[216] | Staple | CAGATGAAAACGAGTAGTAAATTGAAATCTAC |
| 10[166] | 12[167] | Staple | GGCACCAGCCGACAATGACAACACGGTTTATC |
| 7[128] | 7[159] | Staple | AATAACCCCGCCATTACCCAAATCAACGTAAC |
| 0[247] | 2[231] | Staple | GCATTAACATCCAATAAATCATACATAACCTGTTTAGCTATGATAAGAG |
| 12[198] | 12[216] | Staple | CCAAAAGTTGTCGTCTTTCCAGACGTTAGTAAATGAATTGTTGAAAA |
| 1[128] | 1[159] | Staple | TGAGAGTCTGGAGTTTCATTCCATATAACAGT |
| 12[230] | 12[248] | Staple | TTTTCACTTCTGTATGGATTTTGCTAAACAACTTTCAAACTAAAGG |
| 2[38] | 4[39] | Staple | ATAAGCAAAAATTCGCATTAAATGCATCGTAA |
| 10[70] | 12[71] | Staple | GAAGCATGCTGCATTAATGAATCAAGCGGTCC |
| 10[215] | 8[216] | Staple | CATGAGGAATTTGTATCATCGCCTAAAGAGGA |
| 4[119] | 2[120] | Staple | CCCGTCGGTTCCTGTAGCCAGCTTGAATCGAT |
| 0[215] | 2[199] | Staple | CAAAGAATTAGCAAAATTAAGCAATTTGACCATTAGATACGCTTAATTG |
| 10[87] | 8[88] | Staple | ACACAACAGTGGTGCTTGTTACCTGACAGTGC |
| 13[128] | 13[159] | Staple | ACCGTCTATCACGCCTGTAGCATTCCACAGAC |
| 6[230] | 8[231] | Staple | GGAAGAAGGCTTGAGATGGTTTAGAACTGACC |
| 4[166] | 6[167] | Staple | GAATGACGCCAAAAGGAATTACGGAAAGATTC |
| 2[119] | 0[120] | Staple | GAACGGTACTATCAGGTCATTGCCGCGGGAGA |
| 8[55] | 6[56] | Staple | GCACGAATTCTAAGTGGTTGTGAAGCCAGGGT |
| 2[230] | 4[231] | Staple | GTCATTTTAATTCGAGCTTCAAACGTCCAATA |
| 8[198] | 10[199] | Staple | GACCAGGACAAAGTACAACGGAGAGTTTCCAT |
| 10[38] | 12[39] | Staple | TGAGTGACCCGCTTTCCAGTCGGAAAATCCTG |
| 12[119] | 10[120] | Staple | CTGATTGCTGGGCGCCAGGGTGGTAGCTGTTT |

| | | | |
|---|---|---|---|
| 0[87] | 2[71] | Staple | TAGAACCCTCATATATTTTAAATG ATAAATTAATGCCGGACCGGTTGA T |
| 6[166] | 8[167] | Staple | ATCAGTTAAAGCTGCTCATTCAGC CTTCATCA |
| 6[119] | 4[120] | Staple | CTATTACGCAGGCTGCGCAACTG TAGTAACAA |
| 2[198] | 4[199] | Staple | CTGAATAAGGAAGCCCGAAAGAC ATTGAATCC |
| 6[247] | 4[248] | Staple | TACCAGTCCGAGAGGCTTTTGCA ATGTTTAGA |
| 6[159] | 6[128] | Staple | GAGATTTAGGAATACCACATTATC GGTGCGGG |
| 2[166] | 4[167] | Staple | AATATGCGAAGCAAAGCGGATTG AGAAAACGA |
| 8[87] | 6[88] | Staple | GGCCCTGCAAGTGTCCTTAGTGC TGGATGTGC |
| 6[183] | 4[184] | Staple | TACAGGTAAGGCATAGTAAGAGC AATGCTTTA |
| 6[55] | 4[56] | Staple | TTTCCCAGACTCCAGCCAGCTTTC GTTGGTGT |
| 2[87] | 0[88] | Staple | TATGTACCGAGGGTAGCTATTTTT AAAATTTT |
| 9[128] | 9[159] | Staple | TCGAATTCGTAAGAATACACTAAA ACACTCAT |
| 0[183] | 2[167] | Staple | CAGAGCATAAAGCTAAATCGGTT GTGATTCCCAATTCTGCCATGTTT TA |
| 6[70] | 8[71] | Staple | GGGTAACTTCATGCGCACGACTT CATCTGTAA |
| 8[166] | 10[167] | Staple | AGAGTAACTTTGACCCCCAGCGA CACTACGAA |
| 8[38] | 10[39] | Staple | TTGAATCAACTCTGACCTCCTGGG GTGCCTAA |
| 10[198] | 12[199] | Staple | TAAACGGAACCGATATATTCGGTA AAAAGGCT |
| 10[247] | 8[248] | Staple | GCTACAGAATCCGCGACCTGCTC CCAATCATA |
| 4[102] | 6[103] | Staple | GGGAACAAGCGCCATTCGCCATT CCAGCTGGC |
| 12[70] | 12[88] | Staple | ACGCTGGTGTTCCAGTTTGGAAC AAGAGTCCACTATTAACTGAGAGA |
| 6[215] | 4[216] | Staple | GTTAATAACGTTTACCAGACGACG ATCGTCAT |
| 10[119] | 8[120] | Staple | CCTGTGTGTCCCCGGGTACCGAG CTTACGCTC |
| 10[230] | 12[231] | Staple | GACTAAAGAGTTAAAGGCCGCTT ATAATAATT |
| 3[128] | 3[159] | Staple | TCATCAACATTTTACCCTGACTAT TATAGTCA |
| 0[55] | 2[39] | Staple | GAGTAATGTGTAGGTAAAGATTCA ACCATCAATATGATATGAAGATTG T |

| | | | |
|---|---|---|---|
| 10[159] | 10[128] | Staple | ACCTAAAACGAAAGAGGCAAAAT CATGGTCAT |
| 5[128] | 5[159] | Staple | TGGGAAGGGCGCAACTAATGCAG ATACATAAC |
| 4[159] | 4[128] | Staple | CATAAATCAAAAATCAGGTCTAAA TGTGAGCG |
| 8[102] | 10[103] | Staple | TGATACCCGATAAAGACGGAGGA AAATTGTTA |
| 10[183] | 8[184] | Staple | CGTAATGCTTATACCAAGCGCGA ACGCATAGG |
| 0[119] | 2[103] | Staple | AGCCTTTATTTCAACGCAAGGATA GAGAGATCTACAAAGGATCGTAA AA |
| 12[102] | 12[120] | Staple | CCTGGCCAGAACGTGGACTCCAA CGTCAAAGGGCGAAAAGGCAACA G |
| 4[230] | 6[231] | Staple | CTGCGGAATAAAAACCAAAATAGA GGACGTTG |
| 2[247] | 0[248] | Staple | GCTCCTTTTATTTTCATTTGGGGC TAGTAGTA |
| 4[247] | 2[248] | Staple | CTGGATAGGCGAACCAGACCGGA ACTTTAATT |
| 8[70] | 10[71] | Staple | GCAACTCACAGGGCTTAAGCTAC TACGAGCCG |
| 8[247] | 6[248] | Staple | AGGGAACCATTTCAACTTTAATCA GCTCATTA |
| 10[102] | 12[103] | Staple | TCCGCTCGAGGCGGTTTGCGTAT CCTTCACCG |
| 6[87] | 4[88] | Staple | TGCAAGGCGCCGGAAACCAGGCA AAACGGCGG |
| 8[119] | 6[120] | Staple | GCCCTGGAGACAATGTCCCGCCA ACCTCTTCG |
| 11[128] | 11[159] | Staple | TTTTCTTTTCACAGCTTGATACCG ATAGTTGC |
| 8[159] | 8[128] | Staple | TCTTGACAAGAACCGGATATTTTC TAATCTAT |
| 6[102] | 8[103] | Staple | GAAAGGGGAATTGTCAACCTTAT GTGACTCTA |
| 12[38] | 12[56] | Staple | TTTGATGTCAAAAGAATAGCCCGA GATAGGGTTGAGTGTTTTGCCCC |
| 2[215] | 0[216] | Staple | GGCTTAGAATTTCGCAAATGGTCA AGGCAAGG |
| 12[166] | 12[184] | Staple | AGCTTGCAGCCCTCATAGTTAGC GTAACGATCTAAAGTTGAGCCTTT |
| 6[38] | 8[39] | Staple | TGTAAAACCAGGGTGGATGTTCTA TAGGGGCC |
| 8[183] | 6[184] | Staple | CTGGCTGATGAATAAGGCTTGCC CAACATTAT |
| 2[159] | 2[128] | Staple | AACTAAAGTACGGTGTCTGGAAG CAAACAAGA |
| 12[247] | 10[248] | Staple | AATTGCGATTGCGGGATCGTCAC CTAGCAACG |
| 2[183] | 0[184] | Staple | TAGCTCAAGAACGAGTAGATTTAG TAAAGCCT |

| | | | |
|---|---|---|---|
| 8[230] | 10[231] | Staple | AACTTTGGATAAATTGTGTCGAAG GCTTTGAG |
| 12[159] | 12[128] | Staple | TTTCGAGGTGAATTTCTTAAACCA GTGAGACG |
| 2[55] | 0[56] | Staple | AAAAACAGTCAACCGTTCTAGCTG CAATGCCT |
| 0[159] | 0[128] | Staple | TACCAAAAACATTATGACCCTGTA ATACTTTT |
| 2[102] | 4[103] | Staple | CTAGCATTAATTCGCGTCTGGCCA TTCTCCGT |
| 10[55] | 8[56] | Staple | AAGCCTGGTTGGTGTAATGAGTA AGTCGGTGG |
| 2[70] | 4[71] | Staple | AATCAGACATTTTTTAACCAATATA ATGGGAT |
| 12[215] | 10[216] | Staple with elongation for SLB anchoring | CAA GCA GGC GAG CGG ACG AAC TTT TCTCCAAACGCTGAGGCTTGCAG GGACTTTTT |
| 12[55] | 10[56] | Staple with elongation for SLB anchoring | CAA GCA GGC GAG CGG ACG AAC TTT AGCAGGCGGAAACCTGTCGTGCC AAAAGTGTA |
| 4[183] | 2[184] | Staple with elongation for SLB anchoring | CAA GCA GGC GAG CGG ACG AAC TTT AACAGTTCCATCAAAAAGATTAAG TAATGCTG |
| 4[55] | 2[56] | Staple with elongation for SLB anchoring | CAA GCA GGC GAG CGG ACG AAC TTT AGATGGGCTTTTGTTAAATCAGCT AAAGCCCC |
| 12[87] | 10[88] | Staple with elongation for SLB anchoring | CAA GCA GGC GAG CGG ACG AACTTT GTTGCAGCGGCCAACGCGCGGG GAACAATTCC |
| 12[183] | 10[184] | Staple with elongation for SLB anchoring | CAA GCA GGC GAG CGG ACG AAC TTT AATTGTATACCATCGCCCACGCAT GTAAAATA |
| 4[215] | 2[216] | Staple with elongation for SLB anchoring | CAA GCA GGC GAG CGG ACG AAC TTT AAATATTCTTCAAATATCGCGTTTT TGCGGAT |
| 4[87] | 2[88] | Staple with elongation for SLB anchoring | CAA GCA GGC GAG CGG ACG AAC TTT ATTGACCGGGAACGCCATCAAAA AGTCAATCA |

**Table S3. List of staple strands for DOTS in cell-cell experiments**

| Start 5' | End 3' | Note | Sequence |
|---|---|---|---|
| 6[198] | 8[199] | Staple | ACGGAACTGACGAGAAACACCAGCGGTGTACA |
| 8[215] | 6[216] | Staple | CAGATGAAAACGAGTAGTAAATTGAAATCTAC |
| 10[166] | 12[167] | Staple | GGCACCAGCCGACAATGACAACACGGTTTATC |
| 7[128] | 7[159] | Staple | AATAACCCCGCCATTACCCAAATCAACGTAAC |
| 0[247] | 2[231] | Staple | GCATTAACATCCAATAAATCATACATAACCTGTTTAGCTATGATAAGAG |
| 12[198] | 12[216] | Staple | CCAAAAGTTGTCGTCTTTCCAGACGTTAGTAAATGAATTGTTGAAAA |
| 1[128] | 1[159] | Staple | TGAGAGTCTGGAGTTTCATTCCATATAACAGT |
| 10[70] | 12[71] | Staple | GAAGCATGCTGCATTAATGAATCAAGCGGTCC |
| 10[215] | 8[216] | Staple | CATGAGGAATTTGTATCATCGCCTAAAGAGGA |
| 4[119] | 2[120] | Staple | CCCGTCGGTTCCTGTAGCCAGCTTGAATCGAT |
| 0[215] | 2[199] | Staple | CAAAGAATTAGCAAAATTAAGCAATTTGACCATTAGATACGCTTAATTG |
| 10[87] | 8[88] | Staple | ACACAACAGTGGTGCTTGTTACCTGACAGTGC |
| 13[128] | 13[159] | Staple | ACCGTCTATCACGCCTGTAGCATTCCACAGAC |
| 6[230] | 8[231] | Staple | GGAAGAAGGCTTGAGATGGTTTAGAACTGACC |
| 4[166] | 6[167] | Staple | GAATGACGCCAAAAGGAATTACGGAAAGATTC |
| 2[119] | 0[120] | Staple | GAACGGTACTATCAGGTCATTGCCGCGGGAGA |
| 8[55] | 6[56] | Staple | GCACGAATTCTAAGTGGTTGTGAAGCCAGGGT |
| 2[230] | 4[231] | Staple | GTCATTTTAATTCGAGCTTCAAACGTCCAATA |
| 8[198] | 10[199] | Staple | GACCAGGACAAAGTACAACGGAGAGTTTCCAT |
| 12[119] | 10[120] | Staple | CTGATTGCTGGGCGCCAGGGTGGTAGCTGTTT |
| 0[87] | 2[71] | Staple | TAGAACCCTCATATATTTTAAATGATAAATTAATGCCGGACCGGTTGAT |
| 6[166] | 8[167] | Staple | ATCAGTTAAAGCTGCTCATTCAGCCTTCATCA |
| 6[119] | 4[120] | Staple | CTATTACGCAGGCTGCGCAACTGTAGTAACAA |

| | | | |
|---|---|---|---|
| 2[198] | 4[199] | Staple | CTGAATAAGGAAGCCCGAAAGA CATTGAATCC |
| 6[159] | 6[128] | Staple | GAGATTTAGGAATACCACATTAT CGGTGCGGG |
| 2[166] | 4[167] | Staple | AATATGCGAAGCAAAGCGGATTG AGAAAACGA |
| 8[87] | 6[88] | Staple | GGCCCTGCAAGTGTCCTTAGTG CTGGATGTGC |
| 6[183] | 4[184] | Staple | TACAGGTAAGGCATAGTAAGAGC AATGCTTTA |
| 6[55] | 4[56] | Staple | TTTCCCAGACTCCAGCCAGCTTT CGTTGGTGT |
| 2[87] | 0[88] | Staple | TATGTACCGAGGGTAGCTATTTT TAAAATTTT |
| 9[128] | 9[159] | Staple | TCGAATTCGTAAGAATACACTAA AACACTCAT |
| 0[183] | 2[167] | Staple | CAGAGCATAAAGCTAAATCGGTT GTGATTCCCAATTCTGCCATGTT TTA |
| 6[70] | 8[71] | Staple | GGGTAACTTCATGCGCACGACTT CATCTGTAA |
| 8[166] | 10[167] | Staple | AGAGTAACTTTGACCCCCAGCGA CACTACGAA |
| 10[198] | 12[199] | Staple | TAAACGGAACCGATATATTCGGT AAAAAGGCT |
| 4[102] | 6[103] | Staple | GGGAACAAGCGCCATTCGCCAT TCCAGCTGGC |
| 12[70] | 12[88] | Staple | ACGCTGGTGTTCCAGTTTGGAAC AAGAGTCCACTATTAACTGAGAG A |
| 6[215] | 4[216] | Staple | GTTAATAACGTTTACCAGACGAC GATCGTCAT |
| 10[119] | 8[120] | Staple | CCTGTGTGTCCCCGGGTACCGA GCTTACGCTC |
| 10[230] | 12[231] | Staple | GACTAAAGAGTTAAAGGCCGCTT ATAATAATT |
| 3[128] | 3[159] | Staple | TCATCAACATTTTACCCTGACTAT TATAGTCA |
| 10[159] | 10[128] | Staple | ACCTAAAACGAAAGAGGCAAAAT CATGGTCAT |
| 5[128] | 5[159] | Staple | TGGGAAGGGCGCAACTAATGCA GATACATAAC |
| 4[159] | 4[128] | Staple | CATAAATCAAAAATCAGGTCTAA ATGTGAGCG |
| 8[102] | 10[103] | Staple | TGATACCCGATAAAGACGGAGG AAAATTGTTA |
| 10[183] | 8[184] | Staple | CGTAATGCTTATACCAAGCGCGA ACGCATAGG |
| 0[119] | 2[103] | Staple | AGCCTTTATTTCAACGCAAGGAT AGAGAGATCTACAAAGGATCGTA AAA |
| 12[102] | 12[120] | Staple | CCTGGCCAGAACGTGGACTCCA ACGTCAAAGGGCGAAAAGGCAA CAG |

| | | | |
|---|---|---|---|
| 4[230] | 6[231] | Staple | CTGCGGAATAAAAACCAAAATAG AGGACGTTG |
| 8[70] | 10[71] | Staple | GCAACTCACAGGGCTTAAGCTAC TACGAGCCG |
| 6[87] | 4[88] | Staple | TGCAAGGCGCCGGAAACCAGGC AAAACGGCGG |
| 8[119] | 6[120] | Staple | GCCCTGGAGACAATGTCCCGCC AACCTCTTCG |
| 11[128] | 11[159] | Staple | TTTTCTTTTCACAGCTTGATACCG ATAGTTGC |
| 8[159] | 8[128] | Staple | TCTTGACAAGAACCGGATATTTT CTAATCTAT |
| 6[102] | 8[103] | Staple | GAAAGGGGAATTGTCAACCTTAT GTGACTCTA |
| 2[215] | 0[216] | Staple | GGCTTAGAATTTCGCAAATGGTC AAGGCAAGG |
| 12[166] | 12[184] | Staple | AGCTTGCAGCCCTCATAGTTAGC GTAACGATCTAAAGTTGAGCCTT T |
| 8[183] | 6[184] | Staple | CTGGCTGATGAATAAGGCTTGCC CAACATTAT |
| 2[159] | 2[128] | Staple | AACTAAAGTACGGTGTCTGGAAG CAAACAAGA |
| 2[183] | 0[184] | Staple | TAGCTCAAGAACGAGTAGATTTA GTAAAGCCT |
| 8[230] | 10[231] | Staple | AACTTTGGATAAATTGTGTCGAA GGCTTTGAG |
| 12[159] | 12[128] | Staple | TTTCGAGGTGAATTTCTTAAACC AGTGAGACG |
| 2[55] | 0[56] | Staple | AAAAACAGTCAACCGTTCTAGCT GCAATGCCT |
| 0[159] | 0[128] | Staple | TACCAAAAACATTATGACCCTGT AATACTTTT |
| 2[102] | 4[103] | Staple | CTAGCATTAATTCGCGTCTGGCC ATTCTCCGT |
| 10[55] | 8[56] | Staple | AAGCCTGGTTGGTGTAATGAGTA AGTCGGTGG |
| 2[70] | 4[71] | Staple | AATCAGACATTTTTTAACCAATAT AATGGGAT |
| 12[215] | 10[216] | Staple with elongation for cell membrane anchoring | CAA GCA GGC GAG CGG ACG AAC TTT TCTCCAAACGCTGAGGCTTGCA GGGACTTTTT |
| 12[55] | 10[56] | Staple with elongation for cell membrane anchoring | CAA GCA GGC GAG CGG ACG AAC TTT AGCAGGCGGAAACCTGTCGTGC CAAAAGTGTA |
| 4[183] | 2[184] | Staple with elongation for cell membrane anchoring | CAA GCA GGC GAG CGG ACG AAC TTT AACAGTTCCATCAAAAAGATTAA GTAATGCTG |

| | | | |
|---|---|---|---|
| 4[55] | 2[56] | Staple with elongation for cell membrane anchoring | CAA GCA GGC GAG CGG ACG AAC TTT AGATGGGCTTTTGTTAAATCAGC TAAAGCCCC |
| 12[87] | 10[88] | Staple with elongation for cell membrane anchoring | CAA GCA GGC GAG CGG ACG AACTTT GTTGCAGCGGCCAACGCGCGGG GAACAATTCC |
| 12[183] | 10[184] | Staple with elongation for cell membrane anchoring | CAA GCA GGC GAG CGG ACG AAC TTT AATTGTATACCATCGCCCACGCA TGTAAAATA |
| 4[215] | 2[216] | Staple with elongation for cell membrane anchoring | CAA GCA GGC GAG CGG ACG AAC TTT AAATATTCTTCAAATATCGCGTTT TTGCGGAT |
| 4[87] | 2[88] | Staple with elongation for cell membrane anchoring | CAA GCA GGC GAG CGG ACG AAC TTT ATTGACCGGGAACGCCATCAAAA AGTCAATCA |
| 4[255] | 3[255] | Staple with elongation for cell membrane anchoring | CAA GCA GGC GAG CGG ACG AAC TTT TGTTTAGACTGGATAGGCGAACC AGACCGGAA |
| 1[32] | 2[32] | Staple with elongation for cell membrane anchoring | CAA GCA GGC GAG CGG ACG AAC TTT ACCATCAATATGATATGAAGATT GTATAAGCA |
| 3[32] | 4[32] | Staple with elongation for cell membrane anchoring | CAA GCA GGC GAG CGG ACG AAC TTT AAAATTCGCATTAAATGCATCGT AACCGTGCA |
| 9[32] | 10[32] | Staple with elongation for cell membrane anchoring | CAA GCA GGC GAG CGG ACG AAC TTT AACTCTGACCTCCTGGGGTGCC TAATGAGTGA |
| 10[255] | 9[255] | Staple with elongation for cell membrane anchoring | CAA GCA GGC GAG CGG ACG AAC TTT TAGCAACGGCTACAGAATCCGC GACCTGCTCC |
| 2[255] | 1[255] | Staple with elongation for cell membrane anchoring | CAA GCA GGC GAG CGG ACG AAC TTT CTTTAATTGCTCCTTTTATTTTCA TTTGGGGC |
| 12[255] | 11[255] | Staple with elongation for cell membrane anchoring | CAA GCA GGC GAG CGG ACG AAC TTT ACTAAAGGAATTGCGATTGCGG GATCGTCACC |
| 7[32] | 8[32] | Staple with elongation for cell membrane anchoring | CAA GCA GGC GAG CGG ACG AAC TTT CCAGGGTGGATGTTCTATAGGG GCCTTGAATC |

| | | | |
|---|---|---|---|
| 5[32] | 6[32] | Staple with elongation for cell membrane anchoring | CAA GCA GGC GAG CGG ACG AAC TTT GCCTCAGGAAGATCGCTCACGA CGTTGTAAAA |
| 6[255] | 5[255] | Staple with elongation for cell membrane anchoring | CAA GCA GGC GAG CGG ACG AAC TTT GCTCATTATACCAGTCCGAGAGG CTTTTGCAA |
| 11[32] | 12[32] | Staple with elongation for cell membrane anchoring | CAA GCA GGC GAG CGG ACG AAC TTT CCCGCTTTCCAGTCGGAAAATCC TGTTTGATG |
| 8[255] | 7[255] | Staple with elongation for cell membrane anchoring | CAA GCA GGC GAG CGG ACG AAC TTT CAATCATAAGGGAACCATTTCAA CTTTAATCA |

**Supplementary Video 1. Time lapse video showing origami exclusion from the cell spreading area.** A raw 6-min time lapse showing DOTS fluorescence signal change underneath the spreading OT-1 T-cells. In the video, three T-cells landed on the DOTS SLB surface and excluded DOTS from the spreading area resulting in a dark signal. The remaining DOTS clustered and centralized to form cSMAC. Scale bar = 5 μm

**Supplementary Video 2. Single molecule experiments showing the spatiotemporal dynamics of DOTS in the immune synapse**. A raw 5-min time lapse of OT-1 T-cells seeded on low density DOTS SLB surface. Left shows the single molecule DOTS signal and right is the RICM channel showing the T-cell spreading. Scale bar = 5 μm

**Supplementary Video 3. The distribution of F-actin and DOTS at the effector T-cell immune synapse.** Cell spreading (RICM channel), DOTS (red channel), LifeAct-GFP (green channel) were imaged, after 2 min of spreading, for a duration of 20 min. Scale bar = 5 μm

**Supplementary Video 4. 3D view of DOTS and tension patterns at the SSLB-T-cell interface.** The video represents a 360-degree rotation of the SSLB engaging an OT-1 naïve T-cell. Tension signal (gray channel) and DOTS Cy3B signal (green channel) of the SSLB were imaged after 30 min incubation with T-cell. Scale bar = 1 μm
